# A comprehensive annotation and differential expression analysis of short and long non-coding RNAs in 16 bat genomes

**DOI:** 10.1101/738526

**Authors:** Nelly Mostajo Berrospi, Marie Lataretu, Sebastian Krautwurst, Florian Mock, Daniel Desirò, Kevin Lamkiewicz, Maximilian Collatz, Andreas Schoen, Friedemann Weber, Manja Marz, Martin Hölzer

## Abstract

Although bats are increasingly becoming the focus of scientific studies due to their unique properties, these exceptional animals are still among the least studied mammals. Assembly quality and completeness of bat genomes vary a lot and especially non-coding RNA (ncRNA) annotations are incomplete or simply missing. Accordingly, standard bioinformatics pipelines for gene expression analysis often ignore ncRNAs such as microRNAs or long antisense RNAs. The main cause of this problem is the use of incomplete genome annotations. We present a complete screening for ncRNAs within 16 bat genomes. NcRNAs affect a remarkable variety of vital biological functions, including gene expression regulation, RNA processing, RNA interference and, as recently described, regulatory processes in viral infections. Within all investigated bat assemblies we annotated 667 ncRNA families including 162 snoRNAs and 193 miRNAs as well as rRNAs, tRNAs, several snRNAs and IncRNAs, and other structural ncRNA elements. We validated our ncRNA candidates by six RNA-Seq data sets and show significant expression patterns that have never been described before in a bat species on such a large scale. Our annotations will be usable as a resource (Electronic Supplement) for deeper studying of bat evolution, ncRNAs repertoire, gene expression and regulation, ecology, and important host-virus interactions.

**Supplementary information** is available at rna.uni-jena.de/supplements/bats, the Open Science Framework (doi.org/10.17605/OSF.IO/4CMDN), and GitHub (github.com/rnajena/bats_ncrna).

## INTRODUCTION

Bats (Chiroptera) are the most abundant, ecologically diverse and globally distributed animals within all vertebrates^1^, but representative genome arrangements and corresponding coding and non-coding gene annotations are still incomplete^2^. Except for the extreme polar regions, bats can be found throughout the globe, feeding on diverse sources such as insects, blood, nectar, fruits, and pollen^2^. Their origin has been dated in the Cretaceous period, with a diversification explosion process dating back to the Eocene^3^.

The 21 bat families known to date are classified into the suborders *Yinpterochiroptera* and *Yan-gochiroptera*^2,4^ (Fig. 1). Although they account for more than 20 % of the total living mammalian diversity^5^, the genomes of only 16 bat species of the estimated more than 1,300 species^2^ have been sequenced with adequate coverage to date and are publicly available (Fig. 1).

**Figure 1:**
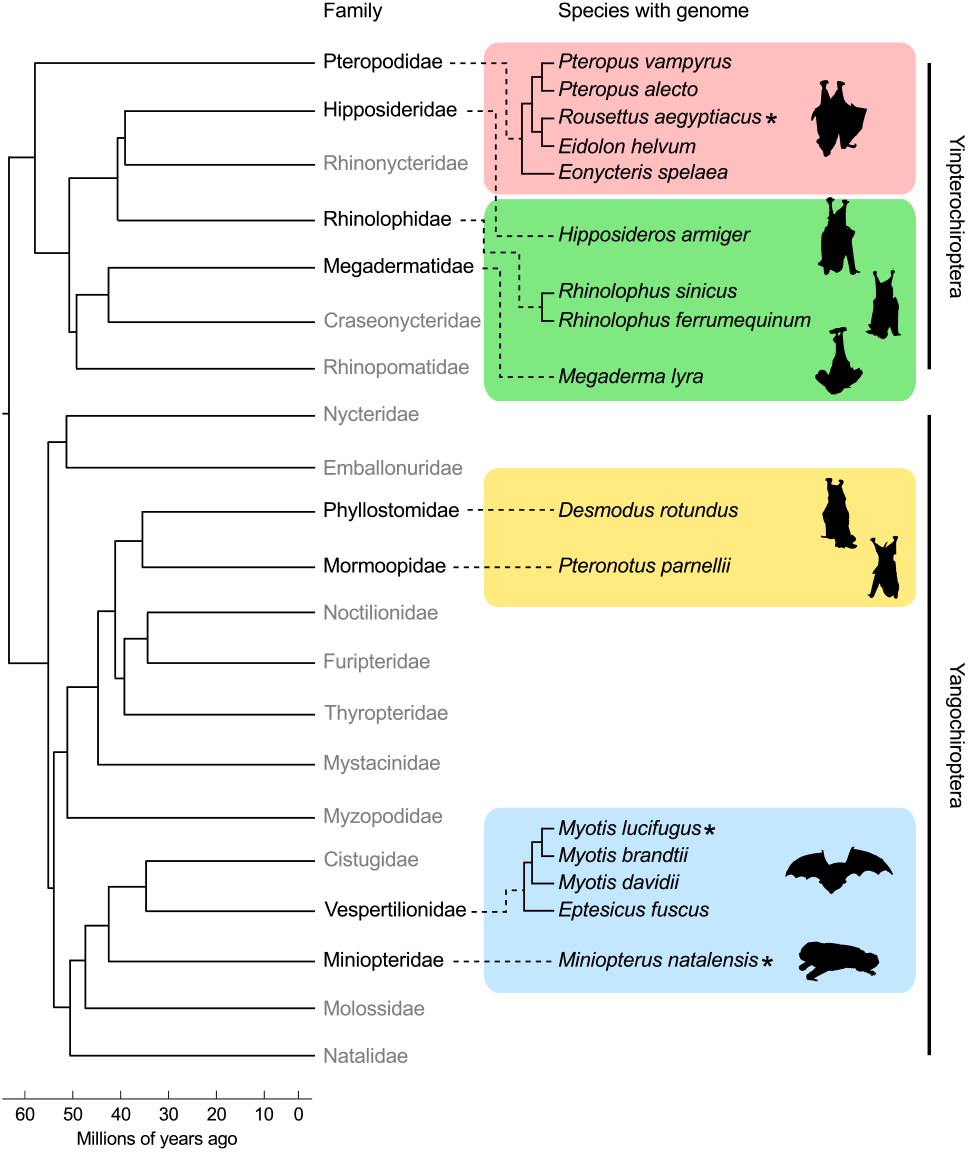
We used available genomes of 16 bat species from eight out of 21 families for non-coding RNA annotation in this study. The tree shows their phylogenetic relationship and is based on a molecular consensus on family relationships of bats ^4^, further adapted and extended from ^2^. Bat families and species with published genomes currently available in the `NCBI` are shown (details see Tab. 1). Bat families still lacking a published genome assembly are written in gray color. RNA-Seq data sets were selected from species marked with an asterisk and additionally obtained from a *Myotis daubentonii* cell line (see Tab. 2). Bat silhouettes were adapted from artworks created by Fiona Reid.

**Table 1:** We have annotated ncRNAs within 16 bat genomes of different assembly quality. We introduced three-letter abbreviations for each bat species used throughout the manuscript and in supplemental files and annotations. Genome sizes were estimated (*Est.*) by using *C*-values (DNA content per pg) from the animal genome size database (http://genomesize.com) and by applying the following formula: *Genome size* = (0.978 · 10^9^) *C*. If multiple entries for one species were available, an average over all *C*-values was calculated and used to estimate the genome size. If one species could not be found, an average *C*-value for the corresponding genus was used. Tab. S1 provides additional assembly statistics calculated by `QUAST` (v5.0.2) ^24^. `NCBI` acc. – GenBank assembly accession without the prefix ‘`GCA`_’.

| Species | Acronym | Size (Gb) | Est. (Gb) | # contigs | N50 | GC (%) | NCBI acc. | Ref. | Year |
| --- | --- | --- | --- | --- | --- | --- | --- | --- | --- |
| *Pteropus vampyrus* | PVA | 2,198 | 2,318 | 36,094 | 5,954,017 | 39.81 | 000151845 | 25 | 2011 |
| *Pteropus alecto* | PAL | 1,985 | 2,025 | 65,598 | 16,439,519 | 39.71 | 000325575 | 26 | 2013 |
| *Rousettus aegyptiacus* | RAE | 1,910 | 2,064 | 2,490 | 2,007,187 | 40.02 | 001466805 | 27 | 2018 |
| *Eidolon helvum* | EHE | 1,837 | 1,985 | 133,538 | 27,684 | 39.24 | 000465285 | 28 | 2013 |
| *Eonycteris spelaea* | ESP | 1,966 | 2,181 | 4,469 | 8,002,591 | 40.30 | 003508835 | 29 | 2018 |
| *Hipposideros armiger* | HAR | 2,236 | 2,516 | 7,571 | 2,328,177 | 41.11 | 001890085 | 30 | 2016 |
| *Rhinolophus sinicus* | RSI | 2,073 | 2,136 | 63,439 | 3,791,306 | 40.51 | 001888835 | 30 | 2016 |
| *Rhinolophus ferrumequinum* | RFE | 1,926 | 2,178 | 160,500 | 21,151 | 40.59 | 000465495 | 28 | 2013 |
| *Megaderma lyra* | MLY | 1,735 | – | 192,872 | 16,881 | 40.39 | 000465345 | 28 | 2013 |
| *Desmodus rotundus* | DRO | 2,063 | 2,519 | 29,801 | 26,869,735 | 42.14 | 002940915 | 31 | 2018 |
| *Pteronotus parnellii* | PPA | 1,960 | 2,533 | 177,401 | 22,675 | 40.82 | 000465405 | 28 | 2013 |
| *Myotis lucifugus* | MLU | 2,034 | 2,210 | 11,654 | 4,293,315 | 42.50 | 000147115 | 25 | 2011 |
| *Myotis brandtii* | MBR | 2,107 | 2,255 | 169,750 | 3,256,877 | 42.55 | 000412655 | 32 | 2013 |
| *Myotis davidii* | MDA | 2,059 | 2,255 | 101,769 | 3,493,173 | 42.69 | 000327345 | 26 | 2013 |
| *Eptesicus fuscus* | EFU | 2,026 | 2,282 | 6,789 | 13,454,942 | 42.93 | 000308155 | – | 2012 |
| *Miniopterus natalensis* | MNA | 1,803 | 1,933 | 1,269 | 4,315,193 | 42.06 | 001595765 | 33 | 2016 |

**Table 2:** Six RNA-Seq data sets comprising alltogether 98 samples derived from four different bat species were used to evaluate our novel ncRNA annotations. All samples were quality trimmed and individually mapped to all 16 bat assemblies using `HISAT2`^34^ and transcript abundances were subsequently calculated from all 1,568 mappings by `featureCounts`^35^. We labeld each RNA-Seq data set based on the first authors last name and the year of data set publication. Raw read data of the enriched sequencing of small RNAs (especially miRNAs) of a *M. daubentonii* cell line (accompanying GSE121301^36^) have been uploaded in the course of this publication under GEO accession GSE132336 (*Weber-2019*). polyA* – library preparation with mRNA selection; rRNA- – library preparation with rRNA depletion and size selection (>200 nt); sRNA – library preparation with size selection (<200 nt); se/pe – single-/paired-end sequencing; ss/not-ss – strand-specific/unstranded sequencing; ss^*s*^/ss^*a*^ – strand-specific in sense orientation/in anti-sense orientation.

| Nr. | Label | Bat species | Nr. samples | Reads |  | Library type | Seq. setup | Accession | Ref. |
| --- | --- | --- | --- | --- | --- | --- | --- | --- | --- |
|  |  |  |  | Length (bp) | Nr. (mio) |  |  |  |  |
| 1 | Field-2015 | *M. lucifugus* | 11 | 101 | 15,1–19,4 | polyA+ | pe/ss<sup>a</sup> | SRP055976 | 37 |
| 2 | Eckalbar-2016 | *M. natalensis* | 18 | 100 | 30,9–82,7 | polyA+ | pe/ss<sup>s</sup> | SRP051253 | 33 |
| 3 | Hölzer-2016 | *R. aegyptiacus* | 9 | 100 | 18,7–25,1 | rRNA- | pe/not-ss | SRP128545 | 16 |
| 4 | Field-2018 | *M. lucifugus* | 24 | 50 | 30,8–49,9 | rRNA- | se/ss<sup>s</sup> | SRP111376 | 38 |
| 5 | Hölzer-2019 | *M. daubentonii* | 18 | 50 | 66,2–72,0 | rRNA- | se/ss<sup>a</sup> | GSE121301 | 36 |
| 6 | Weber-2019 | *M. daubentonii* | 18 | 50 | 8,6–12,5 | sRNA | se/ss<sup>s</sup> | GSE132336 | – |

Bats have developed a variety of unique biological features that are the rarest among all mammalian, including laryngeal echolocation^4,6^, vocal learning^7^, and the ability to fly ^3^. They occupy a broad range of different ecological niches^2^, have an exceptional longevity ^8–10^ and a natural and unique resilience against various pathogenic viruses^1,11^. For example, bats are the suspected reservoirs for some of the deadliest viral diseases such as Ebola and SARS^12–14^, but appear to be asymptomatic and survive the infection. Possibly, the solution to better understand and fight these pathogens lies in the uniquely developed immune system of bats.^15,16^. Studying bats and their genomes is likely to have high impacts on various scientific fields, including healthy ageing, immune and ecosystem functioning, the evolution of sensory perception and human communication, and mammalian genome architecture (see the recent Bat1K review for further details^2^).

Despite the unique biological characteristics of these flying mammals and their important role as natural reservoirs for viruses, bats are one of the least studied taxa of all mammalian^17^. Accordingly, there is little knowledge about the non-protein-coding transcriptome of bats, which plays a crucial role in an extensive number of cellular and regular functions and comprises a very diverse family of untranslated RNA molecules^18,19^. In addition, it is believed that due to the early evolutionary radiation of bats (compared to other mammals) their innate and acquired immune responses have a different set of molecules^1^.

Genome assemblies and annotations are essential starting points for many molecular-driven and comparative studies.^20^. Especially, studies of non-model organisms play important roles in many investigations^21^. In most cases, however, these organisms lack well-annotated genomes^22^, which severely limit our ability to gain a deeper understanding of these species and may further impede biomedical research^23^.

In this study, we comprehensively annotated non-coding RNAs in 16 available bat genome assemblies (Tab. 1). For each bat species, we provide final annotations that are compatible with current `Ensembl` and `NCBI` (National Center for Biotechnology Information) standards (`GTF` format) and that can be directly used in other studies, for example for differential gene expression analysis. We compare our new annotations with the currently available annotations for bats and show that a large number of non-coding genes are simply not annotated and are therefore overlooked by other studies. We used six RNA-Seq data sets comprising different conditions (Tab. 2) to validate our annotations and to determine the expression levels of our newly annotated ncRNAs. Exemplarily, we show that our novel annotations can be used to identify ncRNAs that are significantly differential expressed during viral infections and were missed by previous studies.

## MATERIALS AND METHODS

### Bat species and genomic assembly data

We downloaded the last recent genome versions for 16 bat species in September 2018 from the `NCBI` genome data base (Tab. 1). Within the order of *Yinpterochiroptera*, nine genomic sequences were obtained covering the bat families *Pteropodidae*, *Hipposideridae*, *Rhinolophidae*, and *Megadermatidae* whereas for the order of *Yangochiroptera* another seven genome assemblies were available for the bat families *Phyllostomidae*, *Mormoopidae*, *Vespertilionidae*, and *Miniopteridae* (Fig. 1). We introduced a unique three-letter abbreviation code (Tab. 1) to easily distinguish between the 16 bat species in the manuscript and intermediate annotation files provided in the Electronic Supplement. We used `QUAST` (v5.0.2)^24^ to calculate several assembly statistics for all genomes, shown in Tab. S1.

At the end of 2018, two new bat genomes were presented by the Bat1K project (http://bat1k.ucd.ie/)^2^, comprising a newer version of the greater horseshoe bat genome (*Rhinolophus ferrumequinum*; *Rhinolophidae*) and the genome of the pale spear-nose bat (*Phyllostomus discolor* ; *Phyllostomidae*). However, these two bat genomes were not included in our current study due to the data use policy of the Bat1K consortium and to support a fair and productive use of these data.

### RNA-Seq data

To validate our novel ncRNA predictions, we selected six RNA-Seq data sets^16,33,36–38^ comprising alltogether 98 samples gathered from four different bat species. We have labeled each published RNA-Seq data set based on the first authors last name and the year of data set publication (Tab. 2 and Tab. S2).

All samples were quality trimmed using `Trimmomatic`^39^ (v0.36) with a 4 nt-step sliding-window approach (Q20) and a minimum remaining read length of 20 nt. For the *Field-2018* data set^38^ we additionally removed the three leading 5’ nucleotides from the reads of each sample because of a generally low quality observed by `FastQC` (www.bioinformatics.babraham.ac.uk/projects/fastqc/) (v0.11.7). The remaining reads of all processed samples were individually mapped to all 16 bat assemblies using `HISAT2`^34^ (v2.1.0) and transcript abundances were subsequently calculated from all resulting 1,568 mappings by `featureCounts`^35^ (v1.6.3). If suitable, the appropriate strand-specific counting mode was applied for each data set (see Tab. 2 for information about the strand specificity). For each bat genome assembly, the merged annotation of already known (`NCBI`) and newly identified (this study) ncRNAs was used (Files S1). Due to the size of the annotations and the huge amount of overlaps, long ncRNAs were counted and analyzed for differential expression separately.

To enable a better investigation of small RNAs (sRNAs), we included a data set of the targeted sequencing of sRNAs (especially miRNAs) from *M. daubentonii* cells (*Weber-2019*). To obtain this sRNA sequencing data, total RNA of 18 samples, that was obtained using the same procedure like explained for the rRNA-depleted *M. daubentonii* data^36^, was preprocessed using the Illumina TruSeq smallRNA protocol, sequenced on one HiSeq 2500 lane, and finally uploaded in the course of this study under GEO accession GSE132336. The reads of these 18 sRNA samples were additionally preprocessed by removing potential adapter sequences with `cutadapt`^40^ (v1.8.3) followed by a quality (Q20) trimming using again a window-size of 4 and a minimum length of 15 nt by `PrinSeq`^41^ (v0.20.3). The processed sRNA samples were either individually mapped to the 16 bat genomes for differential expression analysis or combined and mapped on each bat genome to predict known and novel miRNAs with `miRDeep2`^42^.

### Differential gene expression analysis

Only uniquely mapped reads were counted and used for the differential gene expression analyses with `DESeq2`^43^ (v1.16.1). Annotated rRNA genes were removed prior `DESeq2` and TPM (transcripts per million) analysis. All raw read counts from samples of one data set were combined and normalized together using the built-in functionality of DESeq2, followed by pair-wise comparisons to detect significant (adjusted p-value < 0.05; absolute log_2_ fold change > 2) differential expressed ncRNAs.

Besides the `DESeq2` normalization, we calculated TPM values for each ncRNA in each sample as previously described^44^:

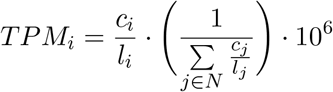

where *c*_*i*_ is the raw read count of ncRNA *i*, *l*_*i*_ is the length of ncRNA *i* (and the cumulative exon length in the case of IncRNAs) and *N* is the number of all ncRNAs in the given annotation. To this end, we obtained for each RNA-Seq sample, each bat annotation, and each ncRNA one TPM value representing the normalized expression level of this ncRNA. If available, we calculated all TPM values in relation to the expression of all already known coding and non-coding genes and not only based on our novel ncRNA annotation.

Although we performed mappings, read countings, and normalization for all samples, bat genome assemblies and all six data sets (Tab. 2; overall 1,568 mappings), we only selected one comparison per data set to exemplarily show novel and significantly differential expressed ncRNAs (Files S2.1-S2.15; divided by data set and input annotation). detected with the help of our extended annotations (Tab. S11). For each data set, we chose the bat species that was also used in the corresponding study. For the *Hölzer-2019* and *Weber-2019* data sets we used the closely related *M. lucifugus* genome assembly and annotation as a reference, because currently no genomic sequence for *M. daubentonii* is publicly available.

#### *Field-2015* (11 samples, MLU)

We compared control (*mock*) samples (5 replicates) with the *infected* (white-nose syndrome, WNS) samples (6 replicates) obtained from wing tissue of *M. lucifugus* to identify novel ncRNAs differentially expressed during the infection with the psychrophilic fungus *Pseudogymnoascus destructans*^37^.

#### *Eckalbar-2016* (18 samples, MNA)

To identify new ncRNAs playing different roles during the development of the bat wing, we compared forelimb samples and hindlimb samples (each of 3 developmental stages in 3 replicates) of *M. natalensis* independent from the embryonic stages^33^.

#### *Hölzer-2016* (9 samples, RAE)

Here, we investigated novel ncRNAs that might play a role due to transcriptional changes between un-infected (*mock*) samples and Ebola/Marburg virus infected samples regardless of the time point of infection and obtained from *R. aegyptiacus* cells^16^. Due to the lack of real biological replicates, we only calculated TPM values and did not include this data set into the differential expression analysis with `DESeq2`.

#### *Field-2018* (24 samples, MLU)

As Field *et al.* were interested in host transcriptomic responses to a fungal pathogen (*P. destructans*, *Pd*) during torpor in hibernating bats (*M. lucifugus*), we searched for novel ncRNAs that were differentially affected between *P. destructans* positive (*Pd+*) samples, obtained although the bats were still torpid (3–6°C; 6 replicates), and *Pd−* samples taken after the bats were allowed to warm to euthermic temperature (6 replicates)^38^.

#### *Hölzer-2019*/*Weber-2019* (18 samples, MLU)

For these studies, RNA was extracted from 18 samples of *M. daubentonii* tissue either left uninfected (*mock*), infected with virus (RVFV Clone 13), or stimulated with interferon (IFN) at two different time points^36^. Sequencing was performed with two different protocols resulting in data sets: *Hölzer-2019* (rRNA-depleted) and *Weber-2019* (smallRNA-enriched). For both data sets we compared *mock* and virus-infected (Clone 13) samples at 24 h post infection (each with 3 replicates) in more detail (Files S2) and present for all 18 smallRNA-Seq samples (*Weber-2019*) normalized expression values in Fig. 4. Please note, that for *M. daubentonii* currently no genome assembly is available, so the genome assembly of *M. lucifugus* was used as a close relative.

### Annotation file format

All of our annotations follow the *General Transfer Format* (`GTF`) as described and defined in the `Ensembl`^45^ data base (https://ensembl.org/info/website/upload/gff.html). Therefore, each row of each annotation file is either defined as a *gene*, *transcript*, or *exon* (by the feature column) and strictly following a hierarchical structure, even if only one exon (as for most ncRNAs) is reported. By adhering to this annotation format, our novel annotations can be easily merged with existing ones as derived from `Ensembl` or `NCBI` and are directly usable as input for computational tools such as `HISAT2` for mapping or `featureCounts` for transcript abundance estimation. We defined gene, transcript, and exon IDs following the `Ensembl` pattern:

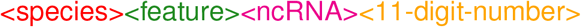

For example, the ID:

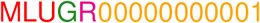

denotes the first (00000000001) rRNA (R) gene (G) annotated in the *M. lucifugs* (MLU) genome. We defined the following abbreviations for different ncRNA types: rRNA (R), tRNA (T), miRNA (M), miRNA with `mirDeep2` (D), snoRNA (S), ncRNA/miscRNA/other (N), IncRNA (L), IncRNA hot-spot (H), mitochondrial ncRNA (O).

### Annotation of non-coding RNAs

In general, we used specialized computational tools for the annotation of specific ncRNA classes (Tab. S3–S10. If not otherwise stated, the main ncRNA-discovery is based on homology searches against the `Rfam` database^46–48^ (v14.0). We used the `Gorap` pipeline (https://github.com/koriege/gorap), a specially developed software suite for genome-wide ncRNA screening. `Gorap` screens genomic sequences for all ncRNAs present in the `Rfam` database using a generalized strategy by applying multiple filters and specialized software tools. To this end, `Gorap` takes huge advantage of `Infernal` (v1.1.2)^49,50^ to annotate ncRNAs based on input alignment files conserved in sequence and secondary structure (so-called *Stockholm* alignment files; `stk`). All resulting alignment files were automatically pre-filtered by `Gorap` based on different ncRNA class-specific parameters including taxonomy, secondary structure, and primary sequence comparisons. Due to repeats, pseudogenes, undiscriminable un-/functional genes, and overlapping results from the different assembly methods, we defined a ncRNA set per species for annotation that includes filtered sequences, but allows for variants and multiple copies. This final annotation set is defined by hand-curating the resulting `stk` alignments of `Gorap` with the help of `Emacs RALEE` mode^51^. Due to the removal of sequences in the stockholm alignments, the remaining sequences were extracted and again aligned into stockholm format using `cmalign -noprob` from `Infernal`. The `Rfam`-derived ncRNA alignments were further split into *snoRNAs* (Tab. S5), *miRNAs* (Tab. S6), and *other ncRNAs* including snRNAs, IncRNAs, and other structural RNAs (Tab. S8).

In general, our annotation results give an overview about the amount of different ncRNAs in bat species and intentionally can include false positive hits and duplicates. All ncRNA hits are placed as `STK` (if available), `GTF` and `FASTA`-files in the Electronic Supplement and OSF (doi.org/10.17605/OSF.IO/4CMDN).

#### rRNAs

We used the prediction tool `RNAmmer` ^52^ (v1.2) to identify 5.8S, 18S and 28S rRNA genes using hidden markov models. The tools output was transformed into regular `GTF` file format. All output files can be found in Tab. S3.

#### tRNAs

For the annotation of tRNAs we applied `tRNAscan-SE`^53^ (v1.3.1) to the bat contigs using default parameters. The results were filtered by removing any tRNAs of type “Undet” or “Pseudo” and the tabular output was transformed into the `GTF` file format. Additional information about the anticodon and the coverage score were added to the description column. We provide the raw `tRNAscan-SE` files in Tab. S4.

#### snoRNAs

We annotated snoRNAs based on available alignments from the `Rfam` data base using `Gorap` and additionally marked and classified them into box C/D and box H/ACA when intersecting with the set of snoRNAs from http://www.bioinf.uni-leipzig.de/publications/supplements/12-022^54^ (Tab. S5).

#### miRNAs

Additionally to the `Rfam`-screening (Tab. S6), miRNAs were predicted by the `miRDeep2` pipeline^42^ (v2.0.0.8) using default parameters and the combined smallRNA-Seq data set (*Weber-2019*; 18 samples) mapped to each individual bat assembly as an input for `miRDeep2` (Tab. S7).

To validate the accuracy of our approach, we compared our `miRDeep2` annotations of *M. lucifugus*/*P. alecto* (based on the transcriptomic data derived from *M. daubentonii*; *Weber-2019*) with annotations of miRNAs for transcriptomic data of *Myotis myotis* ^8^ and *P. alecto*^55^. For reference mapping, Huang *et al.* also used the *M. lucifugus* genome, so we were theoretically able to directly compare our annotations with the annotations of both studies. Unfortunately, no positional information (annotation file) of the identified miRNAs derived from the transcriptomic data of *M. myotis* were given in the manuscript or supplement^8^. Therefore, we blasted the precursor miRNA sequences identified with the help of the *M. myotis* transcriptome against the *M. lucifugus* genome and retained only hits with a sequence identity of 100 %. The so obtained positional information was further used to calculate the overlap between our predicted miRNAs in *M. lucifugus*. We used the same approach for the *P. alecto* comparison. If the annotated location of an miRNA and one of our identified miRNA locations in *M. lucifugus*/*P.alecto* were overlapping by at least 85 %, we counted this location as a common prediction.

#### IncRNAs

Long ncRNAs were annotated using a high confidence data set *H* from the `LNCipedia`^56^ (v5.2) data base comprising 107,039 transcripts of potential human IncRNAs. The transcripts were used as input for a `BLASTn` (2.7.1+, 1*e*^−10^) search against each of the 16 bat assemblies (compiled as `BLAST` databases). The `BLASTn` result for each bat assembly was further processed to group single hits into potential transcripts as follows: first, for each query sequence *q* ∈ *H*, hits of *q* found on the same contig *c* and strand *s* were selected (*hits*_*c*,*s*,*q*_) and the longest one, *q*_1_, was chosen as a starting point, so that *trscp*_*c,s,q*_ = (*q*_1_). Second, all hits *q*_*i*_ ∈ *hits*_*c,s,q*_ with *q*_*i*_ ∉ *trscp*_*c,s,q*_, that do not overlap neither in the query *q* nor in the target sequence and do not exceed a maximum range of 500,000 nt from the most up-stream to the most down-stream target sequence position of all *q*_*j*_ ∈ *trscp*_*c,s,q*_ ⋃ *q*_*i*_, were added iteratively to *trscp*_*c,s,q*_. To this end, we introduced a simple model of exon-intron structures, naturally occurring when using transcript sequences as queries against a target genome assembly. We defined the 500,000 nt search range based on an estimation of IncRNA gene sizes derived from the human `Ensembl`^45^ annotation. If the sum of the lengths of all *q*_*i*_ ∈ *trscp*_*c,s,q*_ covers the query transcript length *length*_*q*_ at least for 70 %, *trscp*_*c,s,q*_ was considered as a transcript and its elements *q*_*i*_ as exons, otherwise all *q*_*i*_ ∈ *trscp*_*c,s,q*_ were withdrawn. This procedure is repeated until all hits ∈ *hits*_*c,s,q*_ were used or withdrawn. Therefore, each so-defined group of non-overlapping hits derived from the same query sequence and found on the same contig and strand should represent a IncRNA transcript with its (rough) exon structure. The defined transcripts were saved as `BLAST`-like output and transformed into `GTF` file format. To follow the `GTF` annotation format and to harmonize our IncRNA annotations with the other ncRNA annotations, each IncRNA transcript was also saved as a gene annotation and consists of at least one exon.

As we observed a lot of different sequences from `LNCipedia` aligning to the same positions in the genomes, we decided to condense exons at the same sequence positions, considering transcripts with one or multiple exons separately. For each contig and strand, starting from the 5’ end, exons with a minimum overlap of 10 nt were grouped together. In the case of multiple exons, groups of exons were merged, if they shared any transcript origin. If all exons in the group originated from the same `LNCipedia` gene, the group was considered as one gene with several transcripts and its associated exon(s). Otherwise, we defined a IncRNA *hot spot* on gene level with several transcripts and their associated exon(s). The `LNCipedia` names of the gathered transcripts of a IncRNA hot spot, as well as start and end positions of all exons, are listed in the `GTF` gene attribute field (Tab. S9). The scripts used for the identification of IncRNAs can be found at https://github.com/rnajena/bats_ncrna.

#### Mitochondrial DNA

As not all of the 16 genome assemblies are including contigs representing the mitochondrial DNA (mtDNA), we downloaded for MLU (KP273591), MBR (KM199849), MDA (KM233172), PAL (NC_023122), PVA (KP214033), PPA (KF752590), RFE (NC_020326), RAE (NC_007393), HAR (NC_018540), and DRO (NC_022423) mitochondrial genomes from the `NCBI` (Tab. 3). For the other six species we used `BLASTn` (2.7.1+, 1*e*^−10^) with the MLU and PVA mitochondrial gemomes as queries against the remaining bat genomes. For EHE we found a possible mtDNA contig in full-length (16,141 nt; AWHC01200796) in the genome assembly. Due to the circularization of mtDNA, we rearranged the sequence of this contig to start with the gene coding for the phenylalanin tRNA and to match the gene order of the other mitochondrial genomes. Only for ESP, RSI, MLY, EFU, and MNA, we were not able to detect any possible contigs of mtDNA (Tab. 3). All eleven mitochondrial genomes were annotated with `MITOS`^57^. The ncRNA results were filtered by e-value (threshold 0.001), thus one of two small rRNAs in MBR and RFE and one of two large rRNAs in EHE were discarded as false positive hits (Tab. S10). For the five bat assemblies directly including mtDNAs (Tab. 3), the MITOS annotations were added to the final merged ncRNA annotation. All other mtDNAs and annotations can be found in Tab. S10.

**Table 3:** Mitochondrial bat genomes (mtDNA) publicly available and used for annotation with `MITOS`^57^. For 10 out of the 16 bat species investigated in this study, mtDNA could be found in the `NCBI`. For four bat species, the mtDNA is also part of the genome assembly as determined using `BLASTn`. For *E. helvum*, no mtDNA could be found in the `NCBI`, but we were able to identify a single contig that is part of the genome assembly as mtDNA using `BLASTn` and the mitochondrial genomes of the other bats as query. The contig was rearranged to match the gene order of the other mtDNAs. ^*R*^ – found via `BLASTn` and rearranged.

| Species | Contig ID in assembly | NCBI accession |
| --- | --- | --- |
| PVA | – | KP214033 |
| PAL | NC_023122 | NC_023122 |
| RAE | NC_007393 | NC_007393 |
| EHE<sup>R</sup> | AWHC01200796 | – |
| ESP | – | – |
| HAR | NC_018540 | NC_018540 |
| RSI | – | – |
| RFE | – | NC_020326 |
| MLY | – | – |
| DRO | NC_022423 | NC_022423 |
| PPA | – | KF752590 |
| MLU | – | KP273591 |
| MBR | – | KM199849 |
| MDA | – | KM233172 |
| EFU | – | – |
| MNA | – | – |

### Computational merging of gene annotations

As we annotated all bat assemblies by using different tools, we needed to merge the resulting `GTF` files to resolve overlapping annotations and to recieve a final annotation of ncRNAs for each bat species. Furthermore, we extended the available `NCBI` annotations (including protein- and non-coding genes) by integrating our novel ncRNA annotations. The scripts used to merge the different annotation files and to calculate overlaps between annotations can be found at https://github.com/rnajena/bats_ncrna. Due to their size, we have not included the IncRNA annotations based on `LNCipedia`. These can be downloaded and used separately (Tab. S9).

#### Merge of novel non-coding annotations

For each bat species we merged the ncRNA annotations (except for IncRNAs) using a custom script (merge_gtf_global_ids.py). After reading in all features and asserting correct file structure, overlaps were resolved in the following manner: (1) exons are considered overlapping if more than 50 % of the shorter one is covered by the larger one. (2) if only one of the overlapping set is of biotype *protein-coding*, remove all others. (3) for further ties, keep only the exon that is highest on a priority list based on annotation source. (4) for further ties, keep only the longest of the exons. For each exon to be removed the corresponding transcript is deleted, and gene records that lost all transcripts are also deleted.

#### Merge of NCBI and novel annotations

We first converted and filtered the `NCBI` annotations to a compatible format with a custom script (format_ncbi.py) and then combined the results with our merged novel ncRNA annotations using the same strategy to resolve overlaps as above, but imposing less strict format rules (merge_gtf_ncbi.py).

## RESULTS

### Assembly and annotation quality differs among bat species

At best, a genome assembly represents the full genetic content of a species at chromosome level. Whereas the first complex eukaryotic genomes were generated using Sanger chemistry, todays technologies such as Illumina short-read sequencing and PacBio or Oxford Nanopore long-read approaches are increasingly used^58^. The currently available bat genomes vary wiedly regarding their assembly quality and completness (Tab. 1, Fig. 2, and Tab. S1) and were predominantly assembled by using short Illumina-derived reads and low (~18 X)^28^ up to moderate/higher coverage (77–218 X) approaches^25,26,30–33^. A new assembly of the cave nectar bat (*Eonycteris spelaea*)^29^ was exclusivly generated from long-read data derived from the PacBio platform and the genome of the Egyptian fruit bat (*Rousettus aegyptiacus*)^27^ was assembled by using a hybrid-approach of Illumina and PacBio data. These two genomes from the *Pteropodidae* family are of a generally higher quality (Fig. 2, Tab. S1). Regardless of their assembly quality, these genomes need to be annotated to identify regions of interest, for example, encoding for protein- and non-coding genes or other regulatory elements.

**Figure 2:**
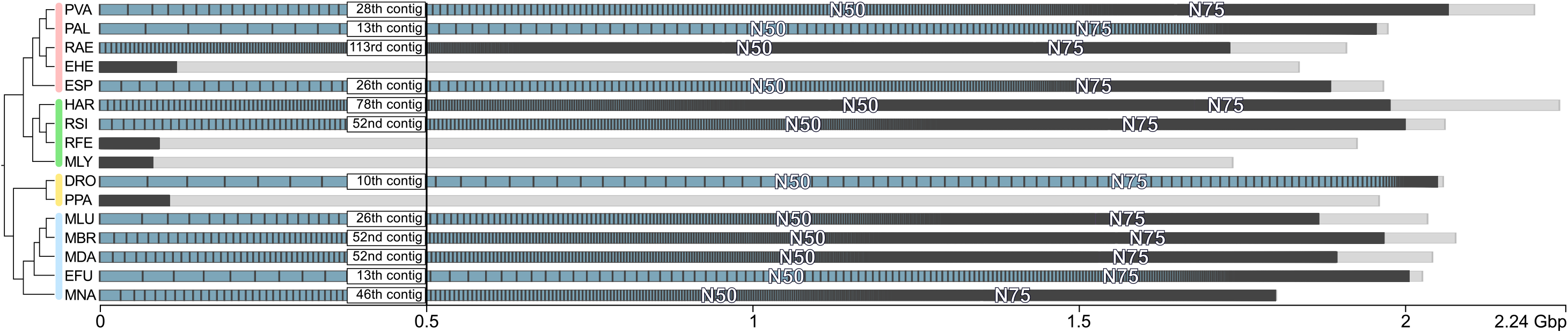
A contig size view of all bat genome assemblies annotated in this study. Only the largest 1,000 contigs of each assembly are shown. Gray bars indicate all contigs that are shorter than the 1,000 largest contigs. N50/N75 labels mark the position of the contig with length equal to the N50/N75 value. Due to a low-coverage approach28, the assemblies of *E. helvum*, *R. ferrumequinum*, *M. lyra*, and *P. parnellii* are of generally lower quality (see Tab. 1 and Tab. S1). Thus, N50/N75 values are not visualized for these species. The figure was adapted from the `Icarus` viewer^59^ used as a part of QUAST^24^.

### Non-coding RNAs are underrepresented in current bat genome annotations

Current genome annotations, mostly generated by automatic annotation pipelines provided by databases such as the `NCBI`^60^ or `Ensembl`^45^, are predominantly focusing on protein-coding genes and well studied ncRNAs such as tRNAs and rRNAs. Accordingly, the available bat genome annotations vary a lot regarding their quality, ranging from more comprehensive annotations for long-standing bat genomes such as *M. lucifugus* or *P. vampyrus* to annotations on *region* level, completely missing any coding or non-coding gene annotations at the current `NCBI` version (Fig. 3 and Tab. 4). Furthermore, and by using strand-specific RNA-Seq data, we could show that some genes (e.g. *IFNA5*/*IFNW2* in the `Ensembl` annotation of *M. lucifugus* ^36^) are annotated on the false strand and are therefore entirely missed by differential expression studies when relying on a strand-specific read quantification. For all publicly available bat genomes, ncRNAs are generally annotated on low levels and are highly incomplete, mostly only comprising some tRNAs, rRNAs, snRNAs, snoRNAs, and IncRNAs (Fig. 3 and Tab. 4). Therefore, many ncRNAs, especially miRNAs, are simply overlooked by current molecular studies, for example from RNA-Seq studies that aim to call differential expressed genes based on such in-complete genome annotation files. Studies that have made additional effort on annotating ncRNAs in bats^8,33,55,61,62^ are not reporting their results on a level that can be directly used for further computational assessment (e.g. as an direct input for RNA-Seq abundance estimation). Currently, in the `NCBI` database, five bat assemblies are entirely lacking any coding/non-coding annotations and miRNAs are not annotated at all (Tab. 4). The `Rfam` database^47^ contains mainly for *M. lucifugus* and *P. vampyrus* 336 ncRNA families. Other ncRNAs are currently unknown from bat genomes or not well documented.

**Table 4:**
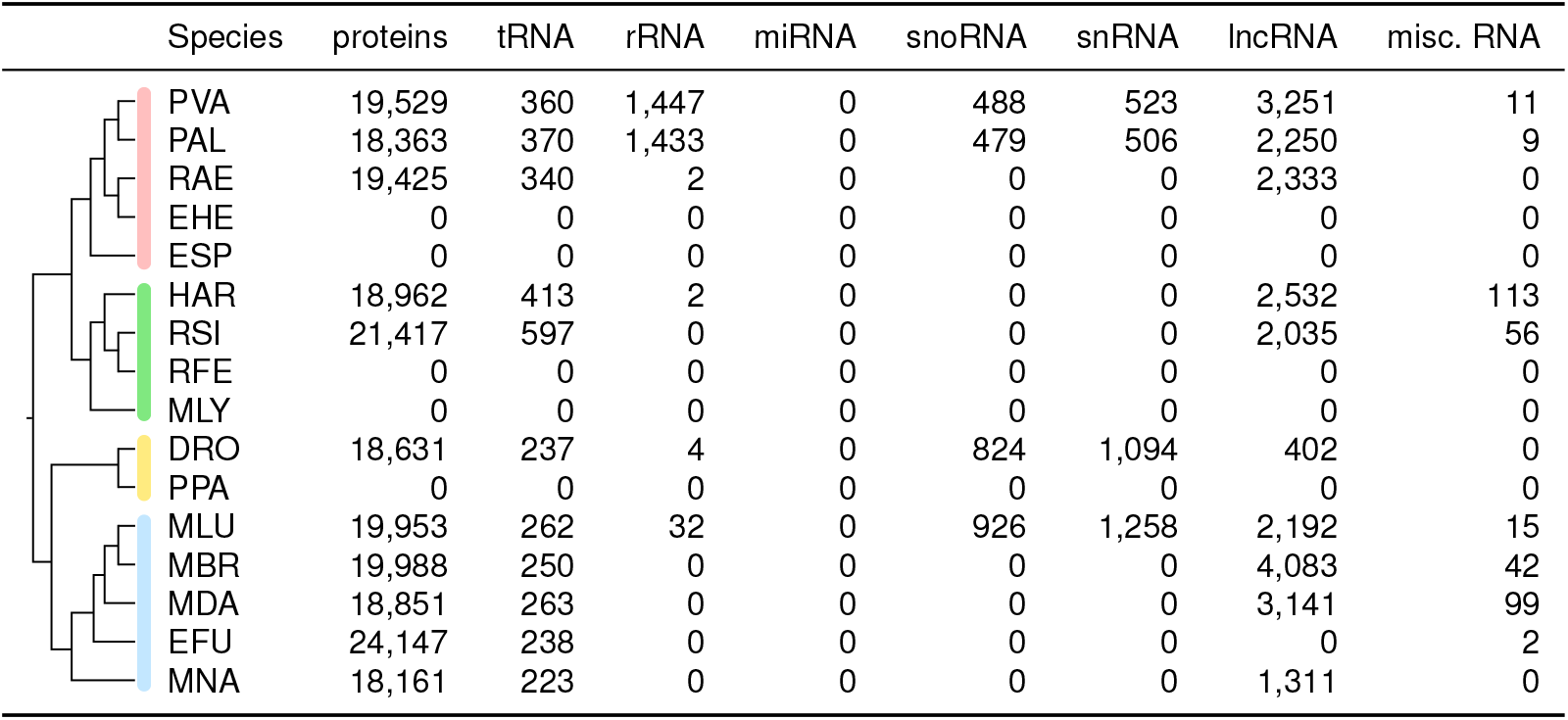
Currently annotated protein-coding genes and ncRNAs of the 16 bat genome assemblies obtained from `NCBI`. Genes were counted by checking the *gene biotype* tag for all *gene* entries (third column in the `GTF` file). Five bat species (EHE, ESP, RFE, MLY, PPA) are currently missing any protein- and non-coding gene annotations and only provide *regions* for each scaffold in the assembly. The annotation of *E. fuscus* is completely missing any *gene biotype* tags. For the description of the three-letter abbreviations please refer to Tab. 1. misc. RNA – miscellaneous RNA, not classifying into the other groups.

| Species | proteins | tRNA | rRNA | miRNA | snoRNA | snRNA | lncRNA | misc. RNA |
| --- | --- | --- | --- | --- | --- | --- | --- | --- |
| PVA | 19,529 | 360 | 1,447 | 0 | 488 | 523 | 3,251 | 11 |
| PAL | 18,363 | 370 | 1,433 | 0 | 479 | 506 | 2,250 | 9 |
| RAE | 19,425 | 340 | 2 | 0 | 0 | 0 | 2,333 | 0 |
| EHE | 0 | 0 | 0 | 0 | 0 | 0 | 0 | 0 |
| ESP | 0 | 0 | 0 | 0 | 0 | 0 | 0 | 0 |
| HAR | 18,962 | 413 | 2 | 0 | 0 | 0 | 2,532 | 113 |
| RSI | 21,417 | 597 | 0 | 0 | 0 | 0 | 2,035 | 56 |
| RFE | 0 | 0 | 0 | 0 | 0 | 0 | 0 | 0 |
| MLY | 0 | 0 | 0 | 0 | 0 | 0 | 0 | 0 |
| DRO | 18,631 | 237 | 4 | 0 | 824 | 1,094 | 402 | 0 |
| PPA | 0 | 0 | 0 | 0 | 0 | 0 | 0 | 0 |
| MLU | 19,953 | 262 | 32 | 0 | 926 | 1,258 | 2,192 | 15 |
| MBR | 19,988 | 250 | 0 | 0 | 0 | 0 | 4,083 | 42 |
| MDA | 18,851 | 263 | 0 | 0 | 0 | 0 | 3,141 | 99 |
| EFU | 24,147 | 238 | 0 | 0 | 0 | 0 | 0 | 2 |
| MNA | 18,161 | 223 | 0 | 0 | 0 | 0 | 1,311 | 0 |

### Identification and validation of ncRNAs in 16 bat genomes

The genome assembly status of different bat species varies widely: ranging for example from 29,801 contigs and an N50 of 26,869,735 nt (*D. rotundus*) to 192,872 contigs and an N50 of only 16,881 nt (*M. lyra*), see Tab. 1 and Tab. S1. Accordingly, also the annotation status varies a lot (Tab. 4). Within this work, we give an overview of potential ncRNA annotations in bats. However, the precise number of ncRNAs remains unclear, because of ncRNAs being present several times in the assemblies, and others still remaining undiscovered.

To give a better estimation of transcribed and potentially functional ncRNAs, we used six Illumina short-read RNA-Seq data sets derived from four bat species (Tab. 2) to estimate the expression levels of our novel annotations. Note that we refer throughout this paper to an RNA-Seq data set by the first author’s name and the year of the respective data set publication. The only included data set derived from a species of the *Yinpterochiroptera* suborder (*R. aegyptiacus*) was obtained from a study dealing with the differential transcriptional responses of Ebola and Marburg virus infections in human and bat cells^16^ (data set: *Hölzer-2016*). In this study, total RNA of nine samples of R06E-J cells, either challenged by the Ebola or Marburg virus or left un-infected, were harvested at 3, 7, or 23 h post infection (poi) and sequenced. Unfortunately, no biological replicates could be generated for this study. Therefore, we did not use this data set for the differential expression analysis, but also calculated normalized expression values (TPM; transcripts per million) as done for all RNA-Seq data sets. The other five data sets comprise *Yangochiroptera* species of the *Vespertilionidae* (*M. lucifugus, M. daubentonii*) and *Miniopteridae* (*M. natalensis*) families (Tab. 2). Field *et al.* conducted two transcriptomic studies^37,38^ (*Field-2015*, *Field-2018*) using wing tissue of the hibernating little brown myotis bat. They were especially interested in transcriptional changes between un-infected wing tissue and adjacent tissue infected with *Pseudogymnoascus destructans*, the fungal pathogen that causes the white-nose syndrome. Two other data sets were obtained from virus-(RVFV Clone 13) and interferon (IFN) alpha-induced transcriptomes of a *Myotis daubentonii* kidney cell line (MyDauNi/2c). RNA of mock, IFN and Clone 13 samples were gathered at two time points, 6 h and 24 h poi^36^. From the same samples, rRNA-depleted (*Hölzer-2019*) and smallRNA-concentrated (*Weber-2019*; see Methods for details) libraries were generated and sequenced. Finally, we used data of the long-fingered bat *M. natalensis* initially obtained to characterize the developing bat wing^33^. Here, total RNA was extracted from paired forelimbs and hindlimbs from three individuals at three developmental stage.

We have mapped each sample to each bat genome, regardless of the origin of the RNA. Expectedly, the mapping rate decreases in bat species that are evolutionary more far away from the original species from which the RNA was sequenced. However, we were interested to find out which ncRNAs are consistently transcribed in all investigated bat species or only in certain bat families and sub-groups.

Over all 16 bat assemblies, we annotated ncRNA families for in total 23 tRNAs, 3 rRNAs, 193 miRNAs (2,680 predicted by `mirDeep2`), 162 snoRNAs, 22 mitochondrial (mt-)tRNAs and 2 mt-rRNAs as well as 244 other ncRNAs additionally derived from the `Rfam` database^47^ (selected ncRNAs are shown in Tab. 5). With a broad approach, we have identified 24,316 potential IncRNAs and defined between 27,149 (*M. lyra*) and 158,135 (*D. rotundus*) IncRNA hot spots.

**Table 5:**
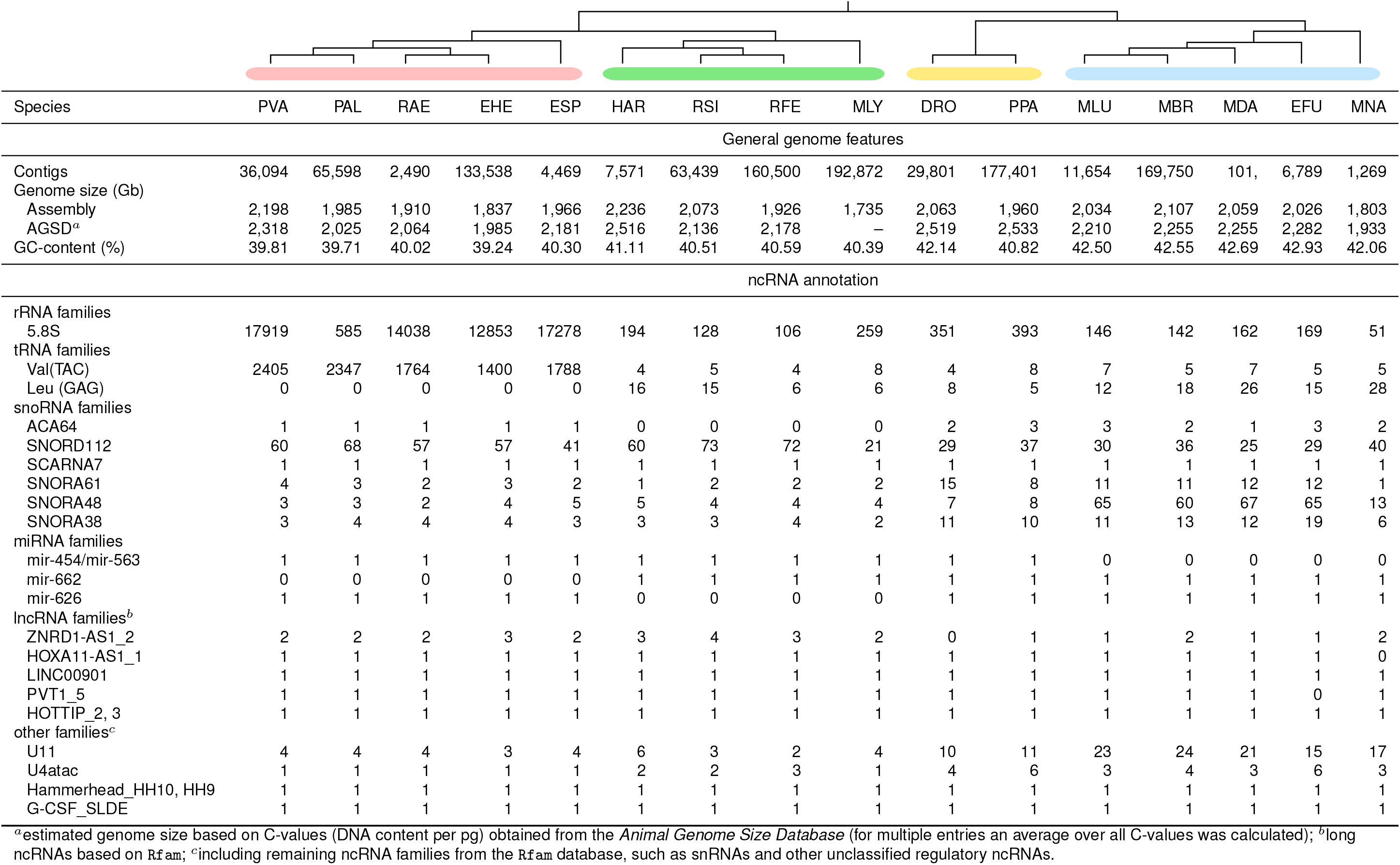
General genome information for each of the 16 investigated bat assemblies and selected ncRNA examples annotated in this study. We selected ncRNAs with interesting copy number distributions among the investigated bat species. For SNORA38, PVT1_5 and HOTTIP_2 and 3 we additionally found sophisticated differential expression patterns in at least one of the used RNA-Seq data sets (absolute log2 fold-change greater 1, TPM> 10). Full tables and detailed information for each ncRNA class (`FASTA, STK, GTF` files) can be found in the Electronic Supplement online (Tab. S3–S10). Members of the *Vespertilionidae* as well as *D. rotundus* and *M. natalensis* appear to have a slightly higher GC-content (+ ~2 %). PVA – *Pteropus vampyrus*; PAL – *Pteropus alecto*; RAE – *Rousettus aegyptiacus*; EHE – *Eidolon helvum*; ESP – *Eonycteris spelaea*; HAR – *Hipposideros armiger* ; RSI – *Rhinolophus sinicus*; RFE – *Rhinolophus ferrumequinum*; MLY – *Megaderma lyra*; DRO – *Desmodus rotundus*; PPA – *Pteronotus parnellii*; MLU – *Myotis lucifugus*; MBR – *Myotis brandtii*; MDA – *Myotis davidii* ; EFU – *Eptesicus fuscus*; MNA – *Miniopterus natalensis*. ACA64 – RF01225; SNORD112 -RF01169; SCARNA7 – RF01295; SNORA61 – RF00420; SNORA48 – RF00554; SNORA38 – RF00428; mir-454 – RF00746; mir-563 – RF01003; mir-662 – RF00983; mir-626 – RF00968; ZNRD1-AS1_2 – RF02219; HOXA11-AS1_1 – RF02137; LINC00901 – RF01884; PVT1_5 – RF02168; HOTTIP_2, 3 – RF02041, RF02042; U11 – RF00548; U4atac – RF00618; Hammerhead_HH10, HH9 – RF02277, RF02275; G-CSF_SLDE – RF00183;

| Species |  |  |  |  |  |  |  |  |  |  |  |  |  |  |  |  |
| --- | --- | --- | --- | --- | --- | --- | --- | --- | --- | --- | --- | --- | --- | --- | --- | --- |
|  | PVA | PAL | RAE | EHE | ESP | HAR | RSI | RFE | MLY | DRO | PPA | MLU | MBR | MDA | EFU | MNA |
| General genome features |  |  |  |  |  |  |  |  |  |  |  |  |  |  |  |  |
| Contigs | 36,094 | 65,598 | 2,490 | 133,538 | 4,469 | 7,571 | 63,439 | 160,500 | 192,872 | 29,801 | 177,401 | 11,654 | 169,750 | 101, | 6,789 | 1,269 |
| Genome size (Gb) |  |  |  |  |  |  |  |  |  |  |  |  |  |  |  |  |
| Assembly | 2,198 | 1,985 | 1,910 | 1,837 | 1,966 | 2,236 | 2,073 | 1,926 | 1,735 | 2,063 | 1,960 | 2,034 | 2,107 | 2,059 | 2,026 | 1,803 |
| AGSD<sup>a</sup> | 2,318 | 2,025 | 2,064 | 1,985 | 2,181 | 2,516 | 2,136 | 2,178 | – | 2,519 | 2,533 | 2,210 | 2,255 | 2,255 | 2,282 | 1,933 |
| GC-content (%) | 39.81 | 39.71 | 40.02 | 39.24 | 40.30 | 41.11 | 40.51 | 40.59 | 40.39 | 42.14 | 40.82 | 42.50 | 42.55 | 42.69 | 42.93 | 42.06 |
| ncRNA annotation |  |  |  |  |  |  |  |  |  |  |  |  |  |  |  |  |
| rRNA families |  |  |  |  |  |  |  |  |  |  |  |  |  |  |  |  |
| 5.8S | 17919 | 585 | 14038 | 12853 | 17278 | 194 | 128 | 106 | 259 | 351 | 393 | 146 | 142 | 162 | 169 | 51 |
| tRNA families |  |  |  |  |  |  |  |  |  |  |  |  |  |  |  |  |
| Val(TAC) | 2405 | 2347 | 1764 | 1400 | 1788 | 4 | 5 | 4 | 8 | 4 | 8 | 7 | 5 | 7 | 5 | 5 |
| Leu (GAG) | 0 | 0 | 0 | 0 | 0 | 16 | 15 | 6 | 6 | 8 | 5 | 12 | 18 | 26 | 15 | 28 |
| snoRNA families |  |  |  |  |  |  |  |  |  |  |  |  |  |  |  |  |
| ACA64 | 1 | 1 | 1 | 1 | 1 | 0 | 0 | 0 | 0 | 2 | 3 | 3 | 2 | 1 | 3 | 2 |
| SNORD112 | 60 | 68 | 57 | 57 | 41 | 60 | 73 | 72 | 21 | 29 | 37 | 30 | 36 | 25 | 29 | 40 |
| SCARNA7 | 1 | 1 | 1 | 1 | 1 | 1 | 1 | 1 | 1 | 1 | 1 | 1 | 1 | 1 | 1 | 1 |
| SNORA61 | 4 | 3 | 2 | 3 | 2 | 1 | 2 | 2 | 2 | 15 | 8 | 11 | 11 | 12 | 12 | 1 |
| SNORA48 | 3 | 3 | 2 | 4 | 5 | 5 | 4 | 4 | 4 | 7 | 8 | 65 | 60 | 67 | 65 | 13 |
| SNORA38 | 3 | 4 | 4 | 4 | 3 | 3 | 3 | 4 | 2 | 11 | 10 | 11 | 13 | 12 | 19 | 6 |
| miRNA families |  |  |  |  |  |  |  |  |  |  |  |  |  |  |  |  |
| mir-454/mir-563 | 1 | 1 | 1 | 1 | 1 | 1 | 1 | 1 | 1 | 1 | 1 | 0 | 0 | 0 | 0 | 0 |
| mir-662 | 0 | 0 | 0 | 0 | 0 | 1 | 1 | 1 | 1 | 1 | 1 | 1 | 1 | 1 | 1 | 1 |
| mir-626 | 1 | 1 | 1 | 1 | 1 | 0 | 0 | 0 | 0 | 1 | 1 | 1 | 1 | 1 | 1 | 1 |
| lncRNA families<sup>b</sup> |  |  |  |  |  |  |  |  |  |  |  |  |  |  |  |  |
| ZNRD1-AS1_2 | 2 | 2 | 2 | 3 | 2 | 3 | 4 | 3 | 2 | 0 | 1 | 1 | 2 | 1 | 1 | 2 |
| HOXA11-AS1_1 | 1 | 1 | 1 | 1 | 1 | 1 | 1 | 1 | 1 | 1 | 1 | 1 | 1 | 1 | 1 | 0 |
| LINC00901 | 1 | 1 | 1 | 1 | 1 | 1 | 1 | 1 | 1 | 1 | 1 | 1 | 1 | 1 | 1 | 1 |
| PVT1_5 | 1 | 1 | 1 | 1 | 1 | 1 | 1 | 1 | 1 | 1 | 1 | 1 | 1 | 1 | 0 | 1 |
| HOTTIP_2, 3 | 1 | 1 | 1 | 1 | 1 | 1 | 1 | 1 | 1 | 1 | 1 | 1 | 1 | 1 | 1 | 1 |
| other families<sup>c</sup> |  |  |  |  |  |  |  |  |  |  |  |  |  |  |  |  |
| U11 | 4 | 4 | 4 | 3 | 4 | 6 | 3 | 2 | 4 | 10 | 11 | 23 | 24 | 21 | 15 | 17 |
| U4atac | 1 | 1 | 1 | 1 | 1 | 2 | 2 | 3 | 1 | 4 | 6 | 3 | 4 | 3 | 6 | 3 |
| Hammerhead_HH10, HH9 | 1 | 1 | 1 | 1 | 1 | 1 | 1 | 1 | 1 | 1 | 1 | 1 | 1 | 1 | 1 | 1 |
| G-CSF_SLDE | 1 | 1 | 1 | 1 | 1 | 1 | 1 | 1 | 1 | 1 | 1 | 1 | 1 | 1 | 1 | 1 |

<sup>a</sup>estimated genome size based on C-values (DNA content per pg) obtained from the *Animal Genome Size Database* (for multiple entries an average over all C-values was calculated); <sup>b</sup>long ncRNAs based on Rfam; <sup>c</sup>including remaining ncRNA families from the Rfam database, such as snRNAs and other unclassified regulatory ncRNAs.

All annotations, separated for each ncRNA class and summarized for each bat species, can be found in the Open Science Frame-work (doi.org/10.17605/OSF.IO/4CMDN) and in our Electronic Supplement (https://www.rna.uni-jena.de/supplements/bats) and are compatible with the genome assembly versions listed in Tab. 1. Thus, our annotations together with the bat genome assemblies obtained from the `NCBI` can be directly used for subsequent analysis such as differential gene expression detection.

### rRNAs

We detected 18S and 28S rRNA for the majority of investigated bat species (Tab. S3). The varying number of rRNAs is in line with all currently available metazoan genomes, lacking the correct composition of rRNAs due to misassemblies. However, the number of 5.8S rRNA varies a lot between 17,919 for *P. vampyrus* and 51 for *M. natalensis* (Tab. 5 and Tab. S3). Interestingly, 4 of 5 *Pteropodidae* show a higher number of 5.8S rRNAs compared to the other species (~50–100 fold). Only the *P. alecto* assembly is in line with the other bat assemblies in regard to the amount of 5.8S rRNA copies.

### tRNAs

For all bat species we observed various numbers of tRNAs (Tab. 5 and Tab. S4). We could detect full sets of tRNAs for *E. fuscus* and *M. davidii*, whereas between one and four tRNAs are missing from the assemblies of *H. armiger*, *M. brandtii*, *M. lucifugus*, *M. lyra*, *M. natalensis*, *P. parnellii*, *R. ferrumequinum*, *R. sinicus*, and *D. rotundus* and between nine and twelve are missing for *E. helvum*, *P. alecto*, *P. vampyrus*, *R. aegyptiacus*, and *E. spelaea* (Tab. S4). Interestingly, we identified a large number of tRNAs (18,235) for *M. lyra* in comparison to all other bat species (Tab. S4), likely a result of the low assembly quality (Fig. 2, Tab. S1). The lowest amount of tRNAs was annotated for *P. parnellii* and *D. rotundus* with only 1,059 and 1,041 copies, respectively. In all other bat genomes we found between 1,735 and 4,657 tRNA genes (Tab. S4).

The tRNA encoding for valine (Val) with the anticodon structure TAC had high copy numbers (over 1,000) in all genome assemblies of the *Pteropodidae* family (Tab. 5). Similar copy numbers were achieved by the *R. ferrumequinum* and *R. sinicus* assemblies regarding the tRNA encoding for isoleucine (Ile) with the anticodon AAT (Tab. S4). For tRNA(Ile) and the anticodon GAT we also observed high copy numbers in *H. armiger*. Interestingly, all species with high tRNA(Val) and tRNA(Ile) copy numbers had rather low counts (between 0 and 28) of tRNA(SeC) with the anticodon TCA, while this tRNA was found with higher copy numbers in *P. parnelli* (68) and in *D. rotundus* (145) and with even higher counts (between 315–498) in all other bat species (Tab. S4). However, high copy numbers might be also occur due to assembly quality and false positive predictions of `tRNAscan-SE`.

### snoRNAs

In Tab. S5 we list all detected snoRNAs, divided into box C/D and box H/ACA types. Overall, we found 162 snoRNA families within the investigated bat species, comprising 88 box C/D, 61 box H/ACA, and 13 unclassified snoRNAs.

Many snoRNAs were found with exactly one copy present in each bat genome assembly (e.g. SCARNA7), whereas others were found in multiple copies for each bat species (e.g. SNORD112) or completely absent from certain bat families (e.g. ACA64), see Tab. 5. Exactly one copy of the small nucleolar RNA ACA64 was found within the genomes of the *Pteropodidae* family and multiple copies for *D. rotundus*, *P. parnellii*, *M. natalensis* and members of the *Vespertilionidae* family, however, this snoRNA seems to be completely absent from bat species of the *Megadermatidae* and *Rhinolophidae* families (Tab. 5). The H/ACA box snoRNA ACA64 is predicted to guide the pseudo-uridylation of 28S rRNA U4331^63^. Interestingly and as another example, SNORA48 was found in higher copy numbers in the genomes of the *Vespertilionidae* family. Among others, this H/ACA box snoRNA was described to be commonly altered in human disease^64^.

### miRNAs

Over all bat assemblies, we detected 193 miRNA families based on `Rfam` alignments (Tab. S6) and predicted between 349 (*E. helvum*) and 2,464 (*M. davidii*) miRNAs based on the 18 combined small RNA-Seq data sets (*Weber-2019*) using `miRDeep2`^42^ (Tab. S7). The higher number of miRNAs predicted for *Myotis* species can be explained because the small RNA-Seq data set is derived from a *Myotis daubentonii* cell line.

Similar to other ncRNA classes, we observe various differences in miRNA copy numbers between the bat families. For example, mir-454 and mir-563 are absent in all *Vespertilionidae* and *M. natalensis* (Tab. 5), whereas mir-1912 is present in all *Yangochiroptera* (except *D. rotundus*) but absent from all *Pteropodidae* (Tab. S6). The miRNA 541 is absent from all *Yangochiroptera* except *D. rotundus*. There are many other examples of absent/present miRNAs in certain bat species/families such as mir-662 (absent from *Pteropodidae*), mir-767 (absent from *Yinpterochiroptera*), and mir-626 (absent from *Rhinolophidae* and *Megaderma lyra*), see Tab. S6.

With `miRDeep2`, we detected hundreds of potential miRNAs for all investigated bat species (Tab. S7). Generally, all 16 bat species can be divided into two groups. For the majority of the bat assemblies (12 out of 16) about 400 miRNAs were predicted. For the other 4 species of the *Vespertilionidae* family about 5 times as many miRNAs could be found. This is in concordance with the small RNA-Seq data used for the prediction, that was obtained from *Myotis daubentonii* kidney cells (see Methods). Nevertheless, ~400 conserved miRNAs can be predicted in the evolutionary more distant bat species.

We compared our `miRDeep2`-predictions in *M. lucifugus* and *P. alecto* with the predictions of two other studies^8,55^. From the 540 published miRNAs based on the transcriptome of *M. myotis* ^8^ and the 426 published miRNAs based on transcriptomic data of *P. alecto*, we were able to obtain 490 and 368 miRNAs with positional information using `BLASTn` against the *M. lucifugus* and *P. alecto* genomes, respectively. From these 490 and 368 miRNAs, our prediction, based on the transcriptome of *M. daubentonii* (*Weber-2019*) mapped to *M. lucifugus* and *P. alecto*, included 195/490 (39.8 %) and 182/368 (49.5 %) miRNAs.

### IncRNAs

For the annotation of IncRNAs, we have deliberately chosen a broad `Blast`-based approach, using 107,039 transcripts of potential human IncRNAs obtained from the `LNCipedia` data base^56^.

We have consciously chosen this approach, because IncRNAs have diverse genomic contexts, reveal various functions and act in different biological mechanisms^65–67^. From 107,039 `LNCipedia` transcripts we annotated 24,316 genes and 182,451 IncRNA *hot spots*. We defined regions in a genome as a IncRNA hot spot, if different `LNCipedia` transcripts derived from different genes map to the same region (see Methods for detailed description). Overall, we found between 58,425 and 137,161 potential IncRNA genes in *M. lucifugus* and *R. sinicus*, respectively, and between 27,149 and 158,135 IncRNA hot spots in *M. lucifugus* and *D. rotundus*, respectively. We annotated the previously described IncRNAs Tbx5-as1 and Hottip ^33^ in all bat genomes, except Tbx5-as1 in *M. lucifugus*, presumably due to the lower genome assembly quality (Tab. 1).

### Other ncRNA elements

Based on the `Rfam` alignments, we were able to detect 244 other ncRNA families in addition to the rRNAs, tRNAs, snoRNAs, and miRNAs described before (Tab. S8). Overlaps with annotated IncRNAs (Tab. S9) are intentional, because the `Rfam` includes only highly structured parts of long ncRNAs.

The highest number of ncRNA copies (951) were detected for the U6 spliceosomal RNA in *M. brandtii*, already known to have a lot of pseudogenes^68^. For 50 ncRNAs such as CAESAR (RF00172), G-CSF_SLIDE (RF00183), NRON (RF00636), TUSC7 (RF01879), Xist_exon4 (RF01881), LINC00901 (RF01884), and Hammerhead_HH9 and _HH10 (RF02275, RF02277) we found exactly one copy in each investigated bat genome assembly (Tab. S8). Again, we also observed ncRNA families that are lost for some species or entire families, for example the ribozyme CoTC (RF00621) that seems to be absent from all *Vespertilionidae* members and *M. natalensis*.

### Mitochondrial annotation

For each investigated bat species except *E. spelaea*, *R. sinicus*, *M. lyra*, *E.fuscus*, and *M. natalensis* (where no mitochondrial contigs could be identified; Tab. 3) mitochondrial protein-coding genes and ncRNAs were annotated (see Methods). In total, 37 mitochondrial genes comprising 22 tRNAs, two rRNAs (12S and 16S) and 13 protein-coding genes were detected for each bat species as known for other metazoans^69^ (Tab. S10). The mitochondrial genome lengths range from 16,343 nt in *E. helvum* to 17,783 nt in *M. davidii*. For the five bat species, where the mitochondrial genome could be identified as a part of the `NCBI` genome assembly, we appended the mtDNA annotation to the final annotation of ncRNAs.

### Updated annotations provide insights into novel differentially expressed ncRNAs

Exemplarily, we investigated known and novel differentially expressed (DE) ncRNAs found in the genome of *M. lucifugus* in more detail. To this end, we used the RNA-Seq data sets *Field-2015*, *Field-2018*, *Hölzer-2019*, and *Weber-2019* (Tab. 2) as a basis to identify DE ncRNAs that were newly discovered in this study and were not part of the current `NCBI` or `Ensembl` genome annotations for this bat species. More detailed DE results can be found in Files S2.

We filtered for novel *M. lucifugus* DE ncRNAs by a) an absolute log_2_ fold change (fc) > 1, b) an adjusted p-value < 0.05, and c) a TPM > 10. We further manually investigated the expression patterns with the `IGV`^71^ and discarded DE ncRNAs overlapping with the current `NCBI` (Myoluc2.0 annotation release 102) or `Ensembl` (Myoluc2.0.96) annotations.

Based on the small RNA-Seq comparison of mock and virus-infected (Clone 13) samples 24 h post infection (*Weber-2019*), we found several miRNAs (`Rfam-` and `mirDeep2`-based) and snoRNAs to be differentially expressed (Fig. 4 and details in Files S2.13). In general, replicates of virus-infected and IFN-treated samples 24 h post infection tend to cluster together only based on the expression profiles of small ncRNAs (mainly miRNAs) (Fig. 4A and B). Most differences can be observed between the 24 h virus-infected and all other samples, that seem to show no clearly distinguishable expression pattern. Interestingly, at 6 h post infection, we see replicates clustering together regardless of their treatment (mock, IFN, Clone 13). Thus, after only 6 h, few miRNAs are differentially expressed and therefore the samples of each replicate (mock, IFN, Clone 13) cluster togehter, because they have the same passaging history but the passaging history in between the replicates differ^36^. After 24 h, more and more miRNAs are significantly differentially expressed and the samples can be better distinguished based on their treatment (Fig. 4A and B). We observed that, in general, miRNAs tend to be down-regulated (Fig 4B; upper half), while snoRNAs tend to be up-regulated (lower half) after 24 h of Clone 13 infection compared to mock. For example, we found a novel miRNA (MLUGD00000002094 in our annotation; predicted by `mirDeep2`; Tab. S7) located in an intron of the protein-coding gene *SEMA3G*, significantly down-regulated (log_2_ fc=−2.56) during Clone 13 infection (Fig. 4C and D). Based on Rfam alignments we further found a histone 3’-UTR stem-loop (RF00032), an RNA element involved in nucleocytoplasmic transport of the histone mRNAs, significantly down-regulated during infection.

**Figure 3:**
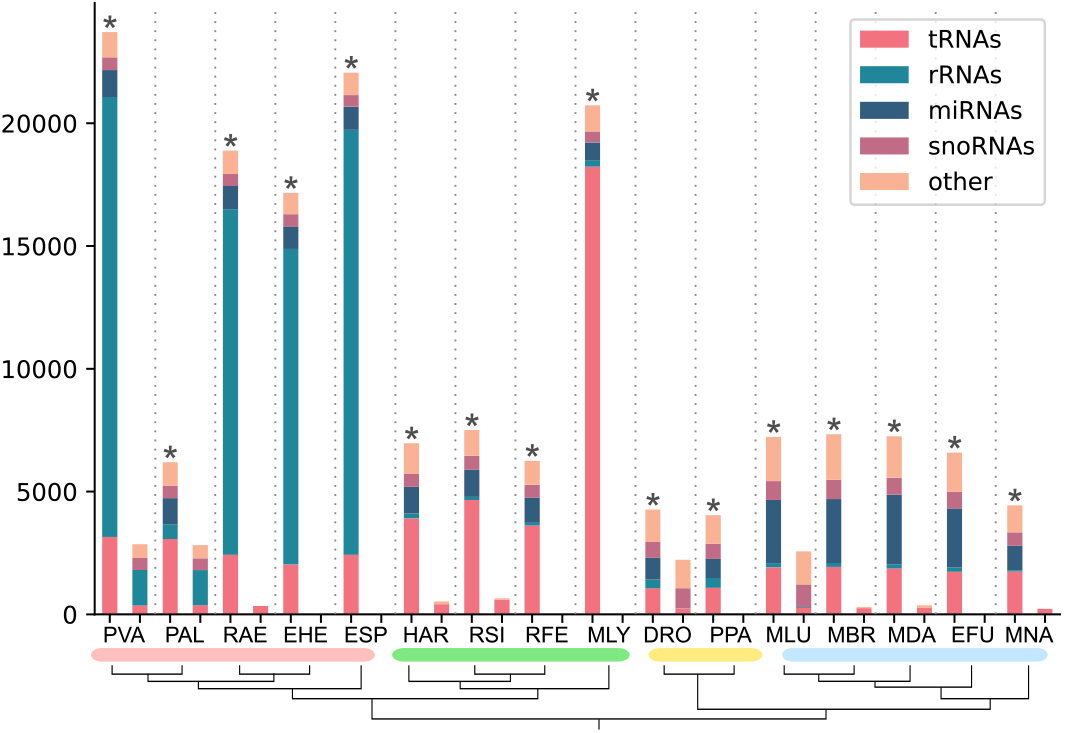
The number of already annotated ncRNAs (`NCBI`) and newly annotated ncRNAs in this study (marked by an asterisk) for each bat species. Due to the general detection approach (see Methods) and their high number IncRNAs are omitted from this figure. Details about IncRNAs can be found in Tab. S9.

**Figure 4:**
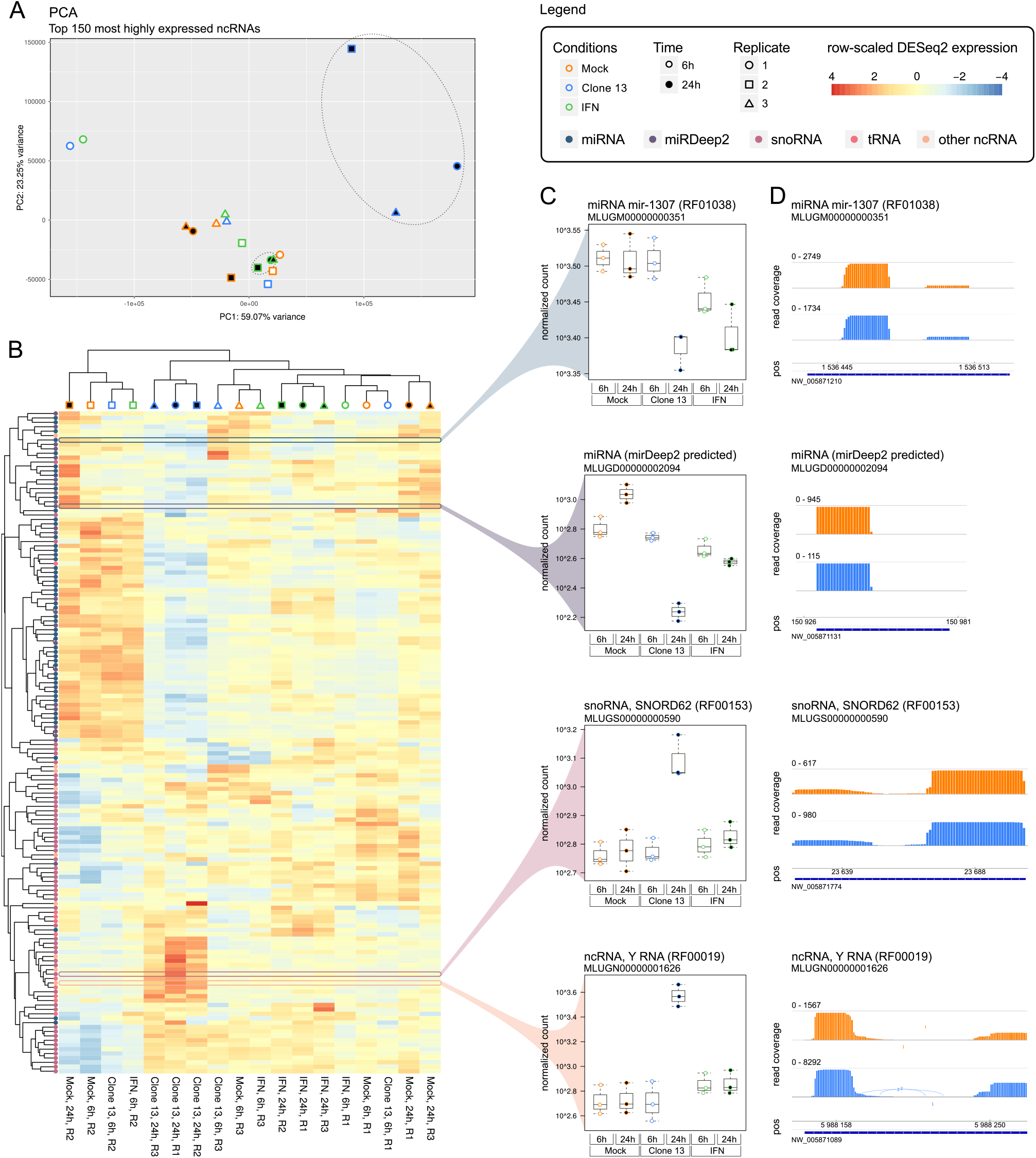
Results shown here are based on expression patterns obtained from the small RNA-Seq samples (*Weber-2019*) mapped to the *M. lucifugus* reference genome (NCBI: GCA_000147115). **(A)** A PCA performed by `PCAGO` ^70^ using the 150 most highly expressed ncRNAs (75 miRNAs, 56 snoRNAs, 10 tRNAs, 9 other small ncRNAs; discarding rRNAs) as input. As also shown in the heat map (B), replicates tend to cluster on the basis of their passaging history, except for virus- and interferon-treated (IFN) samples 24 h post infection. **(B)** A heat map and hirarchical clustering for the same 150 most highly expressed ncRNAs that were used for PCA (A). The clustering was performed on `DESeq2`-normalized read counts and scaled on rows to visualize changes in expression on ncRNA level. Most differences can be observed between the Clone 13-infected samples 24 h post infection and all other conditions. Due to minor changes in miRNA expression levels, the other samples tend to cluster on the basis of their passaging history and not on the basis of their treatment. In general, miRNAs tend to be down-regulated (upper half), while snoRNAs tend to be up-regulated (lower half) after 24 h of Clone 13 infection. **(C)** Selected expression box plots of four highly expressed ncRNAs. Shown are `DESeq2`-normalized expression counts. The miRNA mir-1307 and a novel `mirDeep2`-predicted miRNA are significantly (adjusted p-value < 0.05) down-regulated at 24 h post virus infection. The C/D box snoRNA SNORD62 and a Y RNA are up-regulated. **(D)** `IGV`-derived expression patterns and raw read counts of the four selected ncRNAs obtained from one replicate of the mock and Clone 13 24 h conditions. Blue bars indicate the annotation and strandness.

For the same comparison of mock and virus-infected samples at 24 h, the rRNA-depleted data set (*Hölzer-2019*) revealed several DE IncRNAs. For example, we found a IncRNA potentially transcribed in an intron of *MX1* (MLUGL00000039178) up-regulated (log_2_ fc=3.36) during infection. Another IncRNA (MLUGL00000087396), we found potentially transcribed as a part of two exons of the *PLAT* protein-coding gene and down-regulated (log_2_ fc= −1.67) during viral infection (Files S2.11).

Interestingly, based on the *Field-2015* RNA-Seq data, we found two internal ribosomal entry site (IRES) in the genes *VEGFA* and *ODC* with `Rfam` IDs RF00461 and RF02535, respectively, to be two-fold up-regulated during *P. destructans* infection.

## DISCUSSION

In this study, we comprehensively annotated ncRNAs in 16 readily available bat genomes obtained from the `NCBI` database (Tab. 1). We provide novel annotations in the common `GTF` format, following a hierarchical structure of *gene*, *transcript*, and *exon* features to allow direct integration of our annotations into already available ones (Tab. S3–S10). Finally, we provide for each bat genome assembly an extended annotation file merged with the protein- and non-coding gene annotations that were already available by the `NCBI` database (Files. S1; leaving out potential IncRNAs that can be downloaded separately Tab. S9). We used six RNA-Seq data sets derived from the transcriptomic sequencing of four bat species (Tab. 2) to calculate normalized expression values for our newly annotated ncRNAs and exemplarily show significantly differential expressed ncRNAs (Fig. 4), that were never before described on such a large scale for any bat species.

In addition to the evolutionarily explainable differences in the pure existence and the amount of annotated ncRNAs in bats, we have observed that the assembly quality can also influence the annotation results. While the effects on the annotation of short ncRNAs seem to be small (with some exceptions), the number of identified lncRNAs and lncRNA hot spots increases with increasing assembly quality (e.g. with a higher N50; see Fig. S1). However, this observation may be true for our data and analyses, but it also depends strongly on the annotation method used.

Recently, the Bat1K project (http://bat1k.ucd.ie/) was announced as a global effort to sequence, assemble and annotate high-quality genomes of all living bat species^2^. We aim to extend our annotation of ncRNAs regulary and whenever new bat genomes become publicly available.

In mid January 2019, 16 new low-coverage bat genomes of nine families were submitted by the Broad Institute to the `NCBI` genome database. Unfortunately, our time-consuming and computationally extensive analyses were already completed at this time point. We want to further automate our ncRNA annotation workflow, to easily include these and any new bat (or other mammalian) genomes that will be sequenced and assembled in the future.

Our current identification of ncRNAs in bat species will be usable as a resource (Electronic Supplement) for deeper studying of bat evolution, ncRNAs repertoire, gene expression and regulation, ecology, and important host-virus interactions.

## DATA AVAILABILITY

Detailed information about the bat genomes used in this study, their assembly quality, and all ncRNA candidates (in `FASTA`, `STK`, and `GTF` format) can be found in the Electronic Supplement (rna.uni-jena.de/supplements/bats) Tables S1-S10. The final extended annotations for each investigated bat species can be found in Tab. S11 and the IncRNA annotations in Files S11. To allow full reproducibility of our study, all final and intermediate data files (such as used genome files and mapping files in `BAM` format) were uploaded to the Open Science Framework under accession doi.org/10.17605/OSF.IO/4CMDN. Python scripts used to filter and merge our annotations were deposited at GitHub (github.com/rnajena/bats_ncrna). The virus-infected and IFN-stimulated small RNA-Seq data of the *M. daubentonii* kidney cell line was uploaded to GEO (GSE132336).

## ACKNOWLEDGEMENTS

We thank Ivonne Görlich and Marco Groth from  the Core Facility DNA sequencing of the Leibniz Institute on Aging — Fritz Lipmann Institute in Jena for their help with short-read Illumina sequencing. Martin Hölzer appreciates the support of the Joachim Herz Foundation by the add-on fellowship for inter-disciplinary life science. Kevin Lamkiewicz thanks the Carl Zeiss Foundation for the financial support within the scope of the program line “Breakthroughs”. The authors gratefully acknowledge the help of the developer of the Gorap pipeline, Konstantin Riege.

## FUNDING

This work was supported in part by grants from the *Deutsche Forschungsgemeinschaft* SFB 1021 (B06), SPP 1596 (We 2616/7-1, We 2616/7-2, DR772-10/1, and MA 5082/1-1), MA 5082/9-1, and the LOEWE-Schwerpunkt “Medical RNomics” of the Land Hessen and by the RAPID consortium of the Bundesministerium für Bildung und Forschung (BMBF, grant number 01KI1723E). This study is further part of the Collaborative Research Centre AquaDiva (CRC 1076 AquaDiva) of the Friedrich Schiller University Jena, funded by the Deutsche Forschungsgemeinschaft, as well as the German Centre for Integrative Biodiversity Research (iDiv) Halle-Jena-Leipzig (iDIV FZT 118) of the University of Leipzig, and also supported by the Carl Zeiss Foundation, project “Eine virtuelle Werkstatt für die Digitalisierung in den Wissenschaften” (Durchbrüche).

### Conflict of interest statement

None declared.

## Notes

#### Summary of Updates

Added citation of Figure 3 and changed affiliation of Andreas. Final submission.

https://www.rna.uni-jena.de/supplements/bats

https://doi.org/10.17605/OSF.IO/4CMDN

https://github.com/rnajena/bats_ncrna

## References

[1] C. H. Calisher, J. E. Childs, H. E. Field, K. V. Holmes, and T. Schountz. Bats: important reservoir hosts of emerging viruses. Clin Microbiol Rev, 19(3):531–545, 2006.

[2] E. C. Teeling, S. C. Vernes, L. M. Dávalos, D. A. Ray, M. T. P. Gilbert, E. Myers, and B. Consortium. Bat biology, genomes, and the Bat1K Project: To generate Chromosome-Level genomes for all living bat species. Annu Rev Anim Biosci, (6):23–46, 2018.

[3] N. B. Simmons. An Eocene big bang for bats. Science, 307(5709):527–528, 2005.

[4] E. C. Teeling, G. Jones, and S. J. Rossiter. Phylogeny, genes, and hearing: implications for the evolution of echolocation in bats. In Bat Bioacoustics, pages 25–54. Springer, 2016.

[5] A. Gardner, D. Wilson, and D. Reeder. Mammal Species of the World. A Taxonomic and Geographic Reference. Mammal species of the world: a taxonomic and geographic reference, 12, 2005.

[6] E. C. Teeling, M. S. Springer, O. Madsen, P. Bates, S. J. O’brien, and W. J. Murphy. A molecular phylogeny for bats illuminates biogeography and the fossil record. Science, 307(5709):580–584, 2005.

[7] Y. Prat, L. Azoulay, R. Dor, and Y. Yovel. Crowd vocal learning induces vocal dialects in bats: Playback of conspecifics shapes fundamental frequency usage by pups. PLoS Biol, 15(10):e2002556, 2017.

[8] Z. Huang, D. Jebb, and E. C. Teeling. Blood miRNomes and transcriptomes reveal novel longevity mechanisms in the long-lived bat, *Myotis myotis*. BMC Genom, 17(1):906, 2016.

[9] N. M. Foley, G. M. Hughes, Z. Huang, M. Clarke, D. Jebb, C. V. Whelan, E. J. Petit, F. Touzalin, O. Farcy, G. Jones, et al. Growing old, yet staying young: The role of telomeres in bats’ exceptional longevity. Sci Adv, 4(2):eaao0926, 2018.

[10] Z. Huang, C. V. Whelan, N. M. Foley, D. Jebb, F. Touzalin, E. J. Petit, S. J. Puechmaille, and E. C. Teeling. Longitudinal comparative transcriptomics reveals unique mechanisms underlying extended healthspan in bats. Nat Ecol Evol, 2019.

[11] I. Smith and L.-F. Wang. Bats and their virome: an important source of emerging viruses capable of infecting humans. Curr Opin Virol, 3(1):84–91, 2013.

[12] L.-F. Wang, P. J. Walker, and L. L. Poon. Mass extinctions, biodiversity and mitochondrial function: are bats “special” as reservoirs for emerging viruses? Curr Opin Virol, 1(6):649–657, 2011.

[13] C. E. Brook and A. P. Dobson. Bats as “special” reservoirs for emerging zoonotic pathogens. Trends Microbiol, 23(3):172–180, 2015.

[14] S. J. Anthony, C. K. Johnson, D. J. Greig, S. Kramer, X. Che, H. Wells, A. L. Hicks, D. O. Joly, N. D. Wolfe, P. Daszak, et al. Global patterns in coronavirus diversity. Virus Evol, 3(1), 2017.

[15] T. J. O’shea, P. M. Cryan, A. A. Cunningham, A. R. Fooks, D. T. Hayman, A. D. Luis, A. J. Peel, R. K. Plowright, and J. L. Wood. Bat flight and zoonotic viruses. Emerg Infect Dis, 20(5):741, 2014.

[16] M. Hölzer, V. Krähling, F. Amman, E. Barth, S. H. Bernhart, V. A. O. Carmelo, M. Collatz, G. Doose, F. Eggenhofer, J. Ewald, J. Fallmann, L. M. Feldhahn, M. Fricke, J. Gebauer, A. J. Gruber, F. Hufsky, H. Indrischek, S. Kanton, J. Linde, N. Mostajo, R. Ochsenreiter, K. Riege, L. Rivarola-Duarte, A. H. Sahyoun, S. J. Saunders, S. E. Seemann, A. Tanzer, B. Vogel, S. Wehner, M. T. Wolfinger, R. Backofen, J. Gorodkin, I. Grosse, I. Hofacker, S. Hoffmann, C. Kaleta, P. F. Stadler, S. Becker, and M. Marz. Differential transcriptional responses to Ebola and Marburg virus infection in bat and human cells. Sci Rep, (6):34589, 2016.

[17] A. T. Papenfuss, M. L. Baker, Z.-P. Feng, M. Tachedjian, G. Crameri, C. Cowled, J. Ng, V. Janardhana, H. E. Field, and L.-F. Wang. The immune gene repertoire of an important viral reservoir, the Australian black flying fox. BMC Genom, 13(1):261, 2012.

[18] J. Mattick. Non-coding RNas: the architects of eukaryotic complexity. EMBO Rep, 2(11):986–991, 2001.

[19] O. Isakov, R. Ronen, J. Kovarsky, A. Gabay, I. Gan, S. Modai, and N. Shomron. Novel insight into the non-coding repertoire through deep sequencing analysis. Nucleic Acids Res, 40(11):e86–e86, 2012.

[20] L. Stein. Genome annotation: from sequence to biology. Nat Rev Genet, 2(7):493, 2001.

[21] R. Ekblom and J. Galindo. Applications of next generation sequencing in molecular ecology of non-model organisms. Heredity, 107(1):1, 2011.

[22] S. L. Salzberg. Next-generation genome annotation: we still struggle to get it right. Genome Biology, 20(1): 92, May 2019. ISSN 1474-760X.

[23] W. A. Haynes, A. Tomczak, and P. Khatri. Gene annotation bias impedes biomedical research. Sci Rep, 8(1):1362, 2018.

[24] A. Gurevich, V. Saveliev, N. Vyahhi, and G. Tesler. QUAST: quality assessment tool for genome assemblies. Bioinformatics, 29(8):1072–1075, 2013.

[25] K. Lindblad-Toh, M. Garber, O. Zuk, M. F. Lin, B. J. Parker, S. Washietl, P. Kheradpour, J. Ernst, G. Jordan, E. Mauceli, et al. A high-resolution map of human evolutionary constraint using 29 mammals. Nature, 478(7370):476, 2011.

[26] G. Zhang, C. Cowled, Z. Shi, Z. Huang, K. A. Bishop-Lilly, X. Fang, J. W. Wynne, Z. Xiong, M. L. Baker, W. Zhao, et al. Comparative analysis of bat genomes provides insight into the evolution of flight and immunity. Science, 339(6118):456–460, 2013.

[27] S. S. Pavlovich, S. P. Lovett, G. Koroleva, J. C. Guito, C. E. Arnold, E. R. Nagle, K. Kulcsar, A. Lee, F. Thibaud-Nissen, A. J. Hume, et al. The egyptian rousette genome reveals unexpected features of bat antiviral immunity. Cell, 173(5):1098–1110, 2018.

[28] J. Parker, G. Tsagkogeorga, J. A. Cotton, Y. Liu, P. Provero, E. Stupka, and S. J. Rossiter. Genome-wide signatures of convergent evolution in echolocating mammals. Nature, 502(7470):228, 2013.

[29] M. Wen, J. H. Ng, F. Zhu, Y. T. Chionh, W. N. Chia, I. H. Mendenhall, B. P.-H. Lee, A. T. Irving, and L.-F. Wang. Exploring the genome and transcriptome of the cave nectar bat *Eonycteris spelaea* with PacBio long-read sequencing. GigaScience, 7(10):giy116, 2018.

[30] D. Dong, M. Lei, P. Hua, Y.-H. Pan, S. Mu, G. Zheng, E. Pang, K. Lin, and S. Zhang. The genomes of two bat species with long constant frequency echolocation calls. Mol Biol Evol, page msw231, 2016.

[31] M. L. Z. Mendoza, Z. Xiong, M. Escalera-Zamudio, A. K. Runge, J. Thézé, D. Streicker, H. K. Frank, E. Loza-Rubio, S. Liu, O. A. Ryder, et al. Hologenomic adaptations underlying the evolution of sanguivory in the common vampire bat. Nat Ecol Evol, 2(4):659, 2018.

[32] I. Seim, X. Fang, Z. Xiong, A. V. Lobanov, Z. Huang, S. Ma, Y. Feng, A. A. Turanov, Y. Zhu, T. L. Lenz, et al. Genome analysis reveals insights into physiology and longevity of the Brandt’s bat *Myotis brandtii*. Nat Commun, (4):2212, 2013.

[33] W. L. Eckalbar, S. A. Schlebusch, M. K. Mason, Z. Gill, A. V. Parker, B. M. Booker, S. Nishizaki, C. Muswamba-Nday, E. Terhune, K. A. Nevonen, et al. Transcriptomic and epigenomic characterization of the developing bat wing. Nat Genet, 48(5):528, 2016.

[34] D. Kim, B. Langmead, and S. L. Salzberg. HISAT: a fast spliced aligner with low memory requirements. Nat Methods, 12(4):357–360, 2015.

[35] Y. Liao, G. K. Smyth, and W. Shi. featureCounts: an efficient general purpose program for assigning sequence reads to genomic features. Bioinformatics, 30(7):923–930, 2014.

[36] M. Hölzer, A. Schoen, J. Wulle, M. A. Müller, C. Drosten, M. Marz, and F. Weber. Virus- and interferon alpha-induced transcriptomes of cells from the microbat *Myotis daubentonii*. iScience, 2019.

[37] K. A. Field, J. S. Johnson, T. M. Lilley, S. M. Reeder, E. J. Rogers, M. J. Behr, and D. M. Reeder. The white-nose syndrome transcriptome: activation of anti-fungal host responses in wing tissue of hibernating little brown myotis. PLOS Pathog, 11(10): e1005168, 2015.

[38] K. A. Field, B. J. Sewall, J. M. Prokkola, G. G. Turner, M. F. Gagnon, T. M. Lilley, J. Paul White, J. S. Johnson, C. L. Hauer, and D. M. Reeder. Effect of torpor on host transcriptomic responses to a fungal pathogen in hibernating bats. Mol Ecol, 27(18):3727–3743, 2018.

[39] A. M. Bolger, M. Lohse, and B. Usadel. Trimmomatic: A flexible trimmer for Illumina sequence data. Bioinformatics, (30):2114–2120, 2014. ISSN 1367-4811.

[40] M. Martin. Cutadapt removes adapter sequences from high-throughput sequencing reads. EMBnet.journal, 17(1):10, 2011.

[41] R. Schmieder and R. Edwards. Quality control and preprocessing of metagenomic datasets. Bioinformatics, (27):863–864, 2011. ISSN 1367-4811.

[42] M. R. Friedländer, S. D. Mackowiak, N. Li, W. Chen, and N. Rajewsky. miRDeep2 accurately identifies known and hundreds of novel microRNA genes in seven animal clades. Nucleic Acids Res, (40):37–52, January 2012. ISSN 1362-4962.

[43] M. I. Love, W. Huber, and S. Anders. Moderated estimation of fold change and dispersion for RNA-seq data with DESeq2. Genome Biol, 15(12):550, 2014.

[44] B. Li, V. Ruotti, R. M. Stewart, J. A. Thomson, and C. N. Dewey. RNA-Seq gene expression estimation with read mapping uncertainty. Bioinformatics, (26): 493–500, 2010. ISSN 1367-4811.

[45] D. R. Zerbino, P. Achuthan, W. Akanni, M. R. Amode, D. Barrell, J. Bhai, K. Billis, C. Cummins, A. Gall, C. G. Girón, et al. Ensembl 2018. Nucleic Acids Res, 46 (D1):D754–D761, 2017.

[46] S. Griffiths-Jones, A. Bateman, M. Marshall, A. Khanna, and S. R. Eddy. Rfam: an RNA family database. Nucleic Acids Res, (31):439–441, January 2003. ISSN 1362-4962.

[47] I. Kalvari, J. Argasinska, N. Quinones-Olvera, E. P. Nawrocki, E. Rivas, S. R. Eddy, A. Bateman, R. D. Finn, and A. I. Petrov. Rfam 13.0: shifting to a genome-centric resource for non-coding RNA families. Nucleic Acids Res, 46(D1):D335–D342, 2017.

[48] I. Kalvari, E. P. Nawrocki, J. Argasinska, N. Quinones-Olvera, R. D. Finn, A. Bateman, and A. I. Petrov. Non-coding RNA analysis using the Rfam database. Curr Protoc Bioinformatics, page e51, 2018.

[49] E. P. Nawrocki, D. L. Kolbe, and S. R. Eddy. Infernal 1.0: inference of RNA alignments. Bioinformatics, (25): 1335–1337, 2009.

[50] E. P. Nawrocki and S. R. Eddy. Infernal 1.1: 100-fold faster RNA homology searches. Bioinformatics, (29): 2933–2935, November 2013. ISSN 1367-4811.

[51] S. Griffiths-Jones. RALEE—RNA alignment editor in Emacs. Bioinformatics, (21):257–259, 2005.

[52] K. Lagesen, P. Hallin, E. A. Rødland, H.-H. Staerfeldt, T. Rognes, and D. W. Ussery. RNAmmer: consistent and rapid annotation of ribosomal RNA genes. Nucleic Acids Res, (35):3100–3108, 2007. ISSN 1362-4962.

[53] T. M. Lowe and S. R. Eddy. tRNAscan-SE: a program for improved detection of transfer RNA genes in genomic sequence. Nucleic Acids Res, (25):955–964, 1997.

[54] S. Kehr, S. Bartschat, H. Tafer, P. F. Stadler, and J. Hertel. Matching of Soulmates: coevolution of snoRNAs and their targets. Molec Biol Evol, 31(2): 455–467, 2013.

[55] C. Cowled, C. R. Stewart, V. A. Likic, M. R. Friedländer, M. Tachedjian, K. A. Jenkins, M. L. Tizard, P. Cottee, G. A. Marsh, P. Zhou, et al. Characterisation of novel microRNAs in the Black flying fox (*Pteropus alecto*) by deep sequencing. BMC Genom, 15(1):682, 2014.

[56] P.-J. Volders, J. Anckaert, K. Verheggen, J. Nuytens, L. Martens, P. Mestdagh, and J. Vandesompele. LNCipedia 5: towards a reference set of human long non-coding RNAs. Nucleic Acids Res, (47):D135–D139, January 2019. ISSN 1362-4962.

[57] M. Bernt, A. Donath, F. Jühling, F. Externbrink, C. Florentz, G. Fritzsch, J. Pütz, M. Middendorf, and P. F. Stadler. MITOS: Improved *de novo* metazoan mitochondrial genome annotation. Mol Phylogenet Evol, 69(2):313–319, 2013.

[58] A. R. Khan, M. T. Pervez, M. E. Babar, N. Naveed, and M. Shoaib. A comprehensive study of de novo genome assemblers: current challenges and future prospective. Evol Bioinform, (14):1176934318758650, 2018.

[59] A. Mikheenko, G. Valin, A. Prjibelski, V. Saveliev, and A. Gurevich. Icarus: visualizer for *de novo* assembly evaluation. Bioinformatics, 32(21):3321–3323, 2016.

[60] N. A. O’Leary, M. W. Wright, J. R. Brister, S. Ciufo, D. Haddad, R. McVeigh, B. Rajput, B. Robbertse, B. Smith-White, D. Ako-Adjei, et al. Reference sequence (RefSeq) database at [ncbi: current status, taxonomic expansion, and functional annotation. Nucleic Acids Res, 44(D1):D733–D745, 2015.

[61] T. I. Shaw, A. Srivastava, W.-C. Chou, L. Liu, A. Hawkinson, T. C. Glenn, R. Adams, and T. Schountz. Transcriptome sequencing and annotation for the Jamaican fruit bat (*Artibeus jamaicensis*). PloS One, 7(11):e48472, 2012.

[62] L. Yuan, F. Geiser, B. Lin, H. Sun, J. Chen, and S. Zhang. Down but not out: the role of microRNAs in hibernating bats. PloS One, 10(8):e0135064, 2015.

[63] P. Schattner, S. Barberan-Soler, and T. M. Lowe. A computational screen for mammalian pseudouridylation guide H/ACA RNAs. RNA, 12(1):15–25, 2006.

[64] M. McMahon, A. Contreras, and D. Ruggero. Small RNAs with big implications: new insights into H/ACA snoRNA function and their role in human disease. Wiley Interdisciplinary Reviews: RNA, 6(2):173–189, 2015.

[65] J. T. Y. Kung, D. Colognori, and J. T. Lee. Long noncoding RNAs: past, present, and future. Genetics, (193):651–669, March 2013. ISSN 1943-2631.

[66] J.-W. Nam, S.-W. Choi, and B.-H. You. Incredible RNA: Dual Functions of Coding and Noncoding. Mol Cells, (39):367–374, May 2016. ISSN 0219-1032.

[67] P. Kumari and K. Sampath. cncRNAs: Bi-functional RNAs with protein coding and non-coding functions. Semin Cell Dev Biol, 47-48:40–51, December 2015. ISSN 1096-3634.

[68] M. Marz, T. Kirsten, and P. F. Stadler. Evolution of spliceosomal snRNA genes in metazoan animals. J Mol Evol, 67(6):594–607, 2008.

[69] J. L. Boore. Animal mitochondrial genomes. Nucleic Acids Res, 27(8):1767–1780, 1999.

[70] R. Gerst and M. Hölzer. PCAGO: An interactive web service to analyze RNA-Seq data with principal component analysis. bioRxiv, page 433078, 2018.

[71] H. Thorvaldsdóttir, J. T. Robinson, and J. P. Mesirov. Integrative Genomics Viewer (IGV): high-performance genomics data visualization and exploration. Briefings Bioinf, (14):178–192, 2013. ISSN 1477-4054.

